# The biophysical basis of enterocyte homeostasis

**DOI:** 10.64898/2026.01.28.702213

**Authors:** Peter Hunter, Jarrah Dowrick, Weiwei Ai, David Nickerson, Mohammad Hossein Shafieizadegan, Finbar Argus

## Abstract

We present an approach to analysing cell homeostasis using a ‘bond graph’ modelling approach that ensures that the conservation laws of physics (conservation of mass, charge, and energy, respectively) are satisfied for the interdependent biochemical, electrical, mechanical, and thermal energy storage mechanisms operating within the cell. We apply the bond graph approach to several cell membrane transport mechanisms and then consider how physics constrains intracellular electrolyte homeostasis for enterocytes (the epithelial absorptive cells of the gut). The model includes the electrogenic sodium-potassium ATPase pump (NKA), the glucose transporter (GLUT2), and an inwardly rectifying potassium channel, all in the basolateral membrane, and the electrogenic sodium-driven glucose transporter (SGLT1) in the apical membrane. Glycolysis converts the imported glucose to ATP to drive NKA. For specified levels of sodium, potassium, and glucose in the blood, the model demonstrates how enterocytes absorb sodium and glucose from the gut and transfer glucose to the blood while maintaining the membrane potential and homeostasis of intracellular sodium and potassium. The Gibbs free energy available from the ATP hydrolysis ensures that the cell operates as a ‘sodium battery’ with a high external to internal ratio of sodium concentration in order to provide the energy for many other cellular transport processes. We show that the 3:2 stoichiometry of Na^+^/K^+^ exchange in NKA, coupled with 2:1 Na^+^/glucose cotransport in SGLT1, a 1:2:2 ratio between glucose consumption and ATP and water production in glycolysis, and K^+^ and glucose efflux through Kir and GLUT2, respectively, provides a balanced system that maintains homeostasis of intracellular Na^+^, K^+^, glucose, ATP and water, and homeostasis of the membrane potential, under varying levels of transport of glucose from the gut to the blood. We also show how the flux expressions for SLC transporters, ATPase pumps and ion channels can all be expressed in a consistent and thermodynamically valid way.

## INTRODUCTION

Cellular physiology operates at the theoretical limits set by physical laws – for example, ion channels are sensitive to the passage of a single elementary charge, and retinae can detect one photon [1]. Eukaryotic cells also exploit every form of energy storage mechanism available – biochemical (e.g., solute concentrations and chemical bonds), electrical (e.g., capacitive charge storage in the cell membrane), mechanical (e.g., the elastic compliance of cellular membranes), and thermal (the heat storage essential for maintaining body temperature). The physical processes that maintain intracellular homeostasis are conservation of mass, conservation of charge, and conservation of energy. In this paper we show how the ‘bond graph’ concept, invented over 50 years ago by Henry Paynter at MIT [2] can be used to explain how cellular homeostasis depends on *all* forms of energy transmission, storage, and dissipation (chemical, electrical, mechanical and thermal) via the application of these basic biophysical laws. The application of bond graphs to a variety of biological mechanisms was pioneered by Oster, Perelson and Katchlsky in the 1970s [3] and then later expanded upon by Gawthrop, Crampin, and Pan at the University of Melbourne [4,5,6]. The formulation presented here, including a focus on appropriate units, model reduction strategies, and a new graphical formulation to simplify the analysis, was pioneered by the present authors [7,8].

The application of bond graphs to analysing cellular homeostasis has not, to our knowledge, been undertaken previously and relies on a new algebraic formulation of the steady membrane fluxes [9]. We present a new way of expressing the flux for a reaction that is thermodynamically consistent and applicable across all transmembrane transport mechanisms (exchangers, cotransporters, ATPase pumps and ion channels). We demonstrate the importance of inward rectification in the Kir channel for achieving homeostasis. Other novel contributions in the paper include the demonstration of using bond graphs in whole cell modelling to ensure energy consistency, and an energy-conserving model of an enterocyte for analysis of cell homeostasis.

### UNITS AND THE CONSERVATION LAWS OF PHYSICS

Before we introduce bond graph concepts, it is useful to first discuss units, since adopting a small number of appropriate units and understanding their role in linking atomic-level quantities with processes at the macroscale clarifies the application of bond graphs to multiple physical domains. The SI base units (m, s, kg, mol, A, K, cd) [10] were largely chosen to reflect the technologies available at the time (1960) to measure them with sufficient accuracy (such as the standard kg mass held in a vault in Paris). However, since 2019 the magnitudes of all SI units have been defined by declaring that seven defining constants (the speed of light in vacuum *c* = 2.99792458×10^8^ m.s^-1^, the Planck constant *h* = 6.62607015×10^−34^ J.s, the elementary charge q_e_ = 1.602176634×10^−19^ C, the hyperfine transition frequency of caesium *Δv*_*Cs*_ = 9.192631770×10^9^ s^-1^, the Boltzmann constant k_B_ = 1.380649×10^-23^ J.K^-1^, Avogadro’s constant N_A_ = 6.02214076 ×10^23^ mol^-1^, and the luminous efficacy *K*_*cd*_) have certain exact numerical values when expressed in terms of their SI units [10]. The underlying units used to define these constants are m, s, J, mol, C, K, and cd (the seven SI base units are then defined in terms of these). The first three (m, s, J) establish the energized 4D space-time world in which we reside. The next two (mol, C) reflect atomic or molecular level physical entities: a mole (mol) of substance measures the number of atoms or molecules it contains (there are N_A_ atoms in 12 g of carbon-12); a Coulomb (C) measures charge at the macroscale since the charge on an electron q_e_ scaled up by Avogadro’s number to give Faraday’s constant F= N_A_.q_e_ is the charge in Coulombs per mole (C.mol^-1^). The Kelvin (K) measures the temperature or thermal energy associated with Brownian motion by scaling up the energy per atom or molecule, given by Boltzmann’s constant k_B_ (J.K^-1^), to the macroscale with N_A_.k_B_.T = RT (J.mol^-1^) where R = N_A_.k_B_ is the gas constant. The last unit (Candela or cd) is not needed unless photons need to be counted since a photon of frequency *v* (s^-1^) has energy *hv* (J). Finally, the magnitude of a rotation is expressed in terms of the dimensionless unit radians (rad). The SI base unit kg is J.s^2^.m^-2^, and the SI base unit A is C.s^-1^. From here on, we use these six units: m, s, J, mol, C, and K.

In the following section, we distinguish units that measure the *amount* of a quantity *q*, such as m, mol, and C, from units of 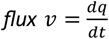 (the amount per second) and potential *u* (expressed as J per unit quantity) driving that flux. We also show that the three conservation laws of physics (conservation of mass, charge, and energy) can be consolidated into a single *conservation of power* law, where power is the product *u. v*.

Bond graphs provide a useful means of formulating and visualising a thermodynamically valid biophysical model because they ensure that mass, charge, and energy are each conserved, and they clearly distinguish the mechanisms for (i) *transmission of power* (the product of potential *u* and flux *v*), (ii) *energy storage* (mechanically in a spring, electrically in a capacitor, or chemically in a solute dissolved in a solvent), (iii) *energy dissipation* to heat (a mechanical damper, an electrical resistance or a chemical reaction), and (iv) *energy transfer* between mechanical, electrical or chemical domains. Most importantly, they distinguish between the conservation laws of physics and the empirically measured constitutive relations that characterise particular materials. For example, a chemical species *i* is stored as a solute 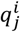 (mol) in a solution at location *j* with a particular *solubility* that generates a chemical potential 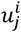 (J.mol^-1^). The diffusion of this solute through a dissipative medium from one location to another is quantified as a molar 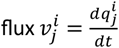 (mol.s^-1^) that depends both on the difference in chemical potential between the two locations and on the *diffusivity* of that medium. The measured values for *solubility* and *diffusivity* are two distinct material constants, and both are quite separate from the equations representing mass and energy conservation. These different material parameters and conservation laws are lumped together in Fick’s law of diffusion.

We use the symbol 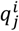 to denote the quantity in moles (mol) of a chemical species *i* at location *j*. Similarly, 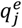 represents electrical charge in Coulombs (C), and 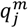 represents distance (m), volume (m^3^) or rotation (rad). Molar flux is denoted 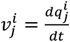 (mol.s^-1^) and 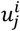 (J.mol^-1^) is the chemical potential generated by storage of solute 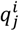 (mol). Electrical current is 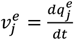 (C.s^-1^) and 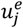 (J.C^-1^) is the electrical potential generated by capacitive storage of charge 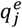 (C). In solid mechanics 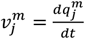 (m.s^-1^) is the velocity and 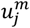 (J.m^-1^) is the force generated by displacement 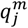 (m), or 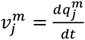 (rad.s^-1^) is the angular velocity and 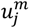 (J.rad^-1^) is the torque generated by rotation 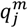 (rad). Finally, in fluid mechanics 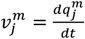 (m^3^.s^-1^) is the fluid flow and 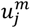 (J.m^-3^) is the pressure (energy density) generated by fluid volume 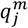 (m^3^).

These units therefore facilitate the expression of *energy flux* (J.s^-1^) as the product of a potential in Joules per unit quantity (mol, C, m, m^3^ or rad) that is driving the flow of that quantity in a way that is common to all physical systems. The product of chemical potential 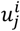 (J.mol^-1^) and flux 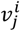 (mol.s^-1^), or electrical potential 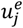 (J.C^-1^) and flux 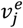 (C.s^-1^), is always power (J.s^-1^). Similarly, the product of mechanical potential (force, pressure, or torque) and mechanical flux (velocity, fluid flow, or angular velocity) is power. The product of heat flow, which is an entropy flux (entropy.s^-1^), and thermal potential (J.entropy^-1^ or temperature in Kelvin K) is also power.

### THE BOND GRAPH MODELLING FRAMEWORK

We now introduce the basic ideas behind the bond graph framework introduced by Paynter [2]. Lines of power transmission called ‘bonds’ always have an associated flux *v* and potential *u* (see Figure 1). If these bonds meet, the sum of powers must be zero: ∑*u. v* = 0, to ensure power conservation. If they share a common potential *u* (called a ‘0:node’), power conservation *u* ∑*v* = 0 becomes just ∑*v* = 0 (for non-zero *u*), which is *mass conservation* if *v* is a molar flux or mechanical flow and *charge conservation* if *v* is an electrical flux. Alternatively, if they share a common flux *v* (called a ‘1:node’), power conservation *v* ∑ *u* = 0 becomes just ∑*u* = 0 (for non-zero *v*), which is *energy conservation*. For chemical reactions these correspond to mass conservation and chemical stoichiometric relations, respectively. For electrical circuits they correspond to Kirchhoff’s current law and voltage law, respectively. For solid mechanics systems they correspond to kinematic consistency and force or torque balance, respectively. For fluid systems they correspond to conservation of volume (conservation of mass which holds when density is assumed constant) and pressure balance.

**Figure 1.**
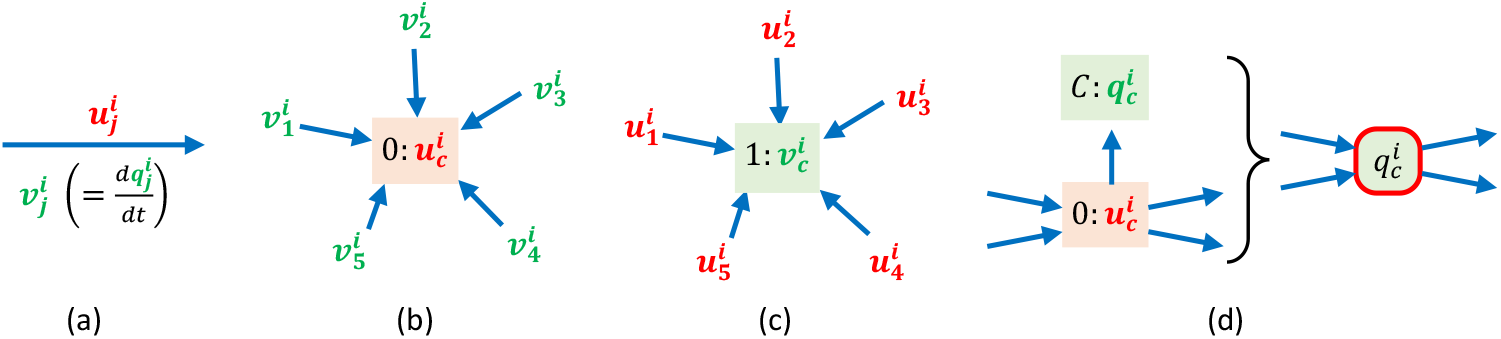
Key bond graph concepts: (a) a bond, which transmits energy, carries a flow 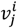 and a potential 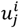; (b) a 0:node is a bond junction where the potential is the same for all bonds and therefore the sum of flows is zero (conservation of mass or charge); (c) a 1:node is a bond junction where the flow is the same for all bonds and therefore the sum of potentials is zero (conservation of energy); (d) a 0:node is usually associated with capacitive energy storage as well as flux balance (top) and can be more succinctly expressed by the red-bordered box where the potential 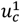 is given by an empirically defined capacitive storage relationship 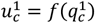. Note that in this figure, the potentials are coloured red and the kinematic quantities in green, just to empathise the difference.

By applying power conservation to 0:nodes and 1:nodes, bond graph models ensure that the conservation laws of physics are always obeyed. By using a common notation (quantity *q*, flux *v*, and potential *u*) with appropriate units (m, s, J, mol, C, K, rad), we can use bond graphs to examine physiological processes that involve the exchange of energy between chemical, electrical, mechanical, and thermal forms.

Note, however, that the mass or charge balance equation ∑ *v* = 0 involves only fluxes *v* (and not potentials *u*), and that the energy balance equation ∑ *u* = 0 involves only potentials *u* (and not flux *v* or quantity *q*). To apply these equations to a specific mechanism, they must therefore be supplemented with *constitutive relations* that couple potential *u* with quantity *q* or flux *v*. Constitutive relations contain the ‘material’ parameters that characterise particular material properties, in contrast with the conservation laws that express the generic constraints imposed by physics. Constitutive laws (and their corresponding material constants) are, for example, needed to describe the static storage mechanisms (solute solubility, electrical charge capacitance, mechanical elasticity, thermal capacitance) and the dissipative mechanisms (chemical reaction kinetics, electrical resistance, mechanical viscosity, and thermal conductivity). Both types of equation are needed but it is extremely important to distinguish the generally applicable conservation laws, captured with 0:nodes and 1:nodes, from the materially specific constitutive equations.

### BOND GRAPH MODELS OF BASIC CELLULAR MECHANISMS

Before considering intracellular homeostasis, we illustrate the application of bond graphs to (i) a simple chemical reaction, (ii) a reaction with multiple reactants and products, (iii) an enzyme-catalysed reaction, (iv) ATP hydrolysis, (v) the glucose transporter (GLUT2), (vi) the electrogenic sodium-glucose cotransporter (SGLT1), (vii) the sodium-potassium ATPase (NKA) pump, and (viii) an inwardly rectifying potassium ion channel (Kir). Wherever experimentally valid, we make assumptions that simplify the models, but always in a way that maintains conservation of mass, charge, and energy. These reduced examples provide the mechanistic bond graph models we subsequently use for analysis of intracellular homeostasis. We also present a simplified graphical representation in order to facilitate clarity as we progress to more advanced models.

#### (i) Simple chemical reaction

We use the simplest possible chemical reaction, consisting of the reversible interconversion of molar quantities 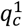 and 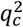 (Figure 2a), to derive the bond graph equations and to discuss the use of bond graph symbols that minimise the complexity of the diagrams.

**Figure 2.**
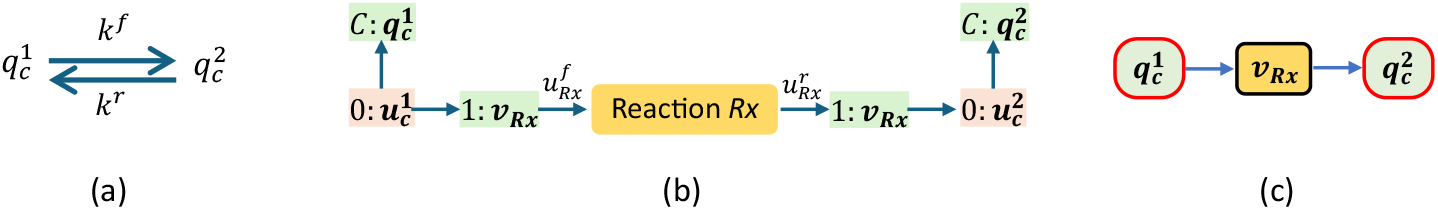
(a) The chemical reaction between 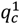 and 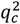; (b) the full bond graph representation; and (c) the simplified bond graph representation in which the red-bordered symbol represents capacitive storage combined with 0:node mass balance, and the black-bordered reaction flux *v*_*Rx*_ symbol includes the reaction and 1:node energy balance. It is sometimes more useful to replace 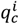 in the red-bordered symbol with the potential 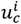 calculated from the 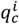 by a Boltzmann equation (see text).

In order to formulate the bond graph describing the simple reaction (Figure 2b), we first identify the molar quantities 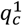 and 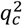 stored in solution as capacitive energy storage elements 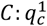 and 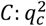, each generating, via a constitutive law (the Boltzmann equation), a chemical potential at a 0:node (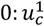 and 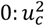) where mass balance is imposed. In Figure 2c the 0:node is combined with the capacitive storage element and written as a single component with a red border (since every molar quantity in solution generates a potential through the Boltzmann equation). In Figure 2b the chemical interconversion between species is a dissipative reaction element interposed between two 1:nodes, enforcing a common reaction flux with associated forward 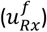 and reverse 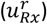 potentials (the arrowhead identifies the positive flux direction). As above, we combine the dissipative element and its neighbouring 1:nodes to form a single simplified element in Figure 2c. In so doing, the seven nodes of the full bond graph representation in Figure 2b are reduced to three in the simplified bond graph (Figure 2c). From here on, we use this simplified representation of the bond graph.

Since the sum of fluxes entering or leaving the first 0:node must be zero:

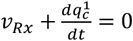

and hence the flux entering the reaction (*v*_*Rx*_) equals the rate of reduction of stored solute 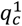:

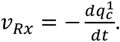

Similarly, on the downstream side (the second 0:node),

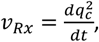

given that the same flux *v*_*Rx*_ passes through the reaction (as highlighted in Figure 2c).

With the bond graph representation in place, we can derive the associated system of equations. If the system is operating at constant temperature and pressure, the Gibbs free energy can be used to characterise the potentials on either side of the reaction, and for a dilute system the chemical potential (the Gibbs free energy per mole) is given by the Boltzmann thermodynamic relation [12],

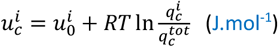

where 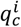 is the number of moles of chemical species 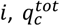 is the total number of moles of all substances in the mixture in compartment 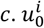 is the (reference) potential when 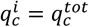.

More compactly,

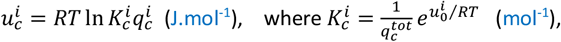

is the constitutive law for biochemical energy storage, and 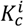 (mol^-1^) is a thermodynamic parameter. To simplify the equations, we introduce the non-dimensional quantity

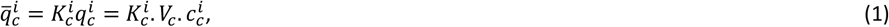

where 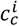 is the concentration of chemical species *i, V*_*c*_ is the volume of *c*.

The chemical potential is then

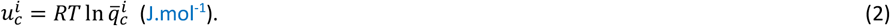

Energy balance at the 1:nodes is

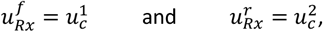

where 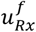 and 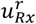 are the forward and reverse potentials for reaction *Rx*. In this case, with only one chemical species entering the reaction, the energy balance is trivial, but if there were multiple reactants and/or multiple products, the 1:node energy balance ensures the appropriate chemical stoichiometry for the reaction.

The molar flow for the reaction *Rx* can be given by the Marcelin-de Donder formula [4] (a constitutive relation but one that, like the Boltzmann relation, can be derived from assumptions about the distribution of particle velocities):

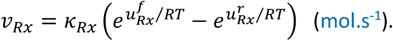

where *κ*_*Rx*_ (mol.s^-1^) is the experimentally determined reaction rate constant (a constitutive parameter).

Substituting for the potentials gives

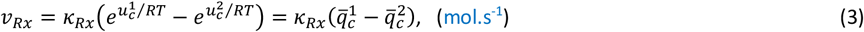

with equilibrium (zero flow) achieved when 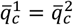.

The change in Gibbs free energy per mole for the reaction is 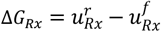, which must be negative for the reaction to proceed (the second law of thermodynamics), with heat output rate 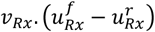. Note that heat output rate being the potential difference times the flow is the same in a biochemical reaction as it is in an electrical resistance or a mechanical damper but the flux for a biochemical reaction depends on the difference in the *exponentials* of 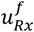 and 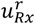 (the Marcelin-de Donder formula above), whereas the flux through an electrical resistor or mechanical damper is dependent only on the potential difference across the resistor or damper (the voltage drop or net force, respectively). At equilibrium, the change in Gibbs free energy is zero.

Note that, from now on, every reaction is assumed to have associated potentials 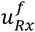 and 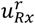 driving the forward and reverse reactions, respectively, and these will not be explicitly labelled on the bond graph diagram.

In summary, the potentials 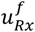 and 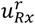 that drive the reaction in the forward and reverse directions are obtained (via energy balance relations) from the chemical solute potentials 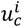 that depend (via Boltzmann’s constitutive equation) on the nondimensional terms 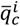 (which include the thermodynamic constants 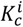). The nondimensional solute quantity 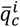 is expressed in terms of the solute concentration 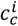 (in compartment c with volume *V*_*c*_) by 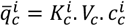. The change in Gibbs free energy occurring in the reaction is 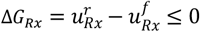 (Δ*G*_*Rx*_ = 0 at equilibrium). The reaction flux *v*_*Rx*_ is given (via the Marcelin-de Donder formula) by equation (3) and the heat output is −Δ*G*_*Rx*_. *v* _*Rx*_

#### (ii) A reaction with multiple reactants and products

Most reactions involve multiple reactants and products, often with variable stoichiometry. These can all be represented as illustrated in Figure 3 (including multiple instances of one chemical species, where, for example, 2 moles of one species combine with 1 mole of another species).

**Figure 3.**
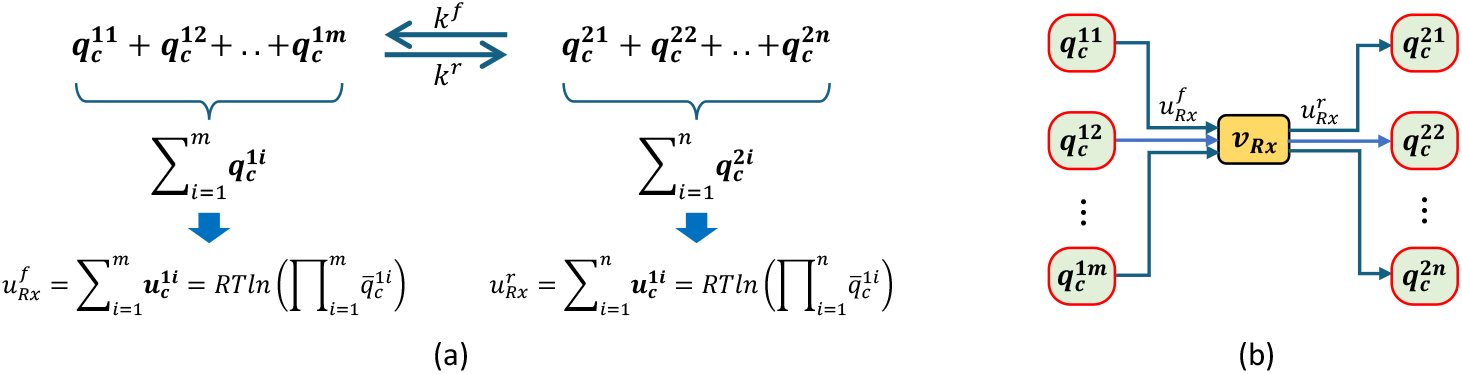
(a) A reaction with multiple reactants and products. 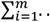 and 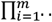 imply a sum and product, respectively, over the *m* reactants, and 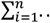 and 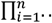 imply a sum and product over the *n* reaction products. (b) The corresponding bond graph diagram.

The reaction flux is

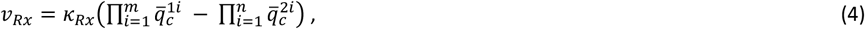

and the change in Gibbs free energy is

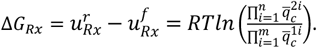

In terms of solute concentrations (using equation 1),

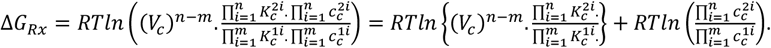

This can be written

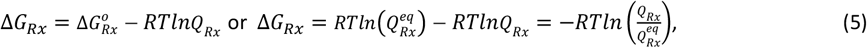

where

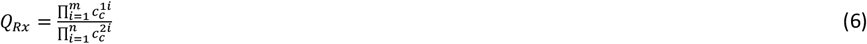

is the ratio of reactant concentrations to product concentrations,

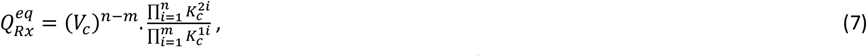

is the equilibrium constant (i.e., when 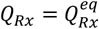, then 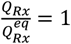 and Δ*G*_*Rx*_ = 0), and

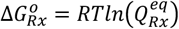

is the free energy needed to move the system from the reference state (all quantities at the reference concentration of 1M and hence *Q*_*Rx*_ = 1 or *RT. lnQ*_*Rx*_ = 0) to the equilibrium state where 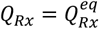 and Δ*G*_*Rx*_ = 0.

Note that the equilibrium constant 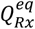, obtained from the measured ratio of reactant concentrations to product concentrations at equilibrium (when *v*_*Rx*_ and Δ*G*_*Rx*_ are zero), along with the reaction rate (see below), is an important parameter characterising a reaction. It has a value of 1 for a simple diffusive process, since the flow is zero when the reactant and product concentrations are equal. When the flow *v*_*Rx*_ is non-zero, the ratio 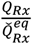 (with a corresponding 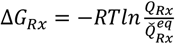) characterizes how far the reaction is from equilibrium under those conditions.

Summarising, the change in Gibbs free energy (the *free energy of the reaction*) is

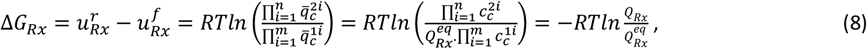

a relationship we will make much use of below. The thermodynamic constants for each chemical species have been combined into just one constant 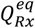, whose value can be established simply by measuring the ratio 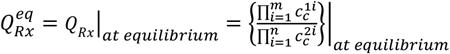. For a reaction to proceed in the forward direction requires Δ*G*_*Rx*_ < 0 or 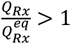 (the second law of thermodynamics).

Note that the reaction flux given by (4) can also be expressed in terms of concentrations:

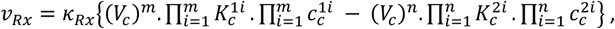

or

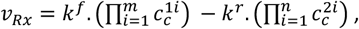

where 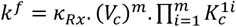 and 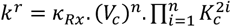.

This is the “mass action” equation widely used in the biochemical literature but note that the parameters *k*^*f*^ and *k*^*r*^ combine two completely different material quantities: the *thermodynamic* Boltzmann coefficient 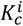 associated with capacitive storage of solute *i* (which generates the chemical potential 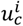 via the empirical Boltzmann equation), and the *reaction rate constant κ*_*Rx*_, which is a material property of the reaction.

A preferable way of representing the reaction is to define

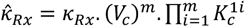

(which depends on both the actual reaction rate and the thermodynamic constants of the reactants) and then

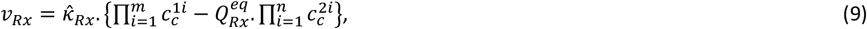

highlighting the fact that only two parameters are needed to describe the reaction, one that can be obtained from steady state concentrations, and one that specifies how quickly the reaction occurs.

We illustrate these concepts with a simple sodium-chloride reaction in Figure 4.

**Figure 4.**
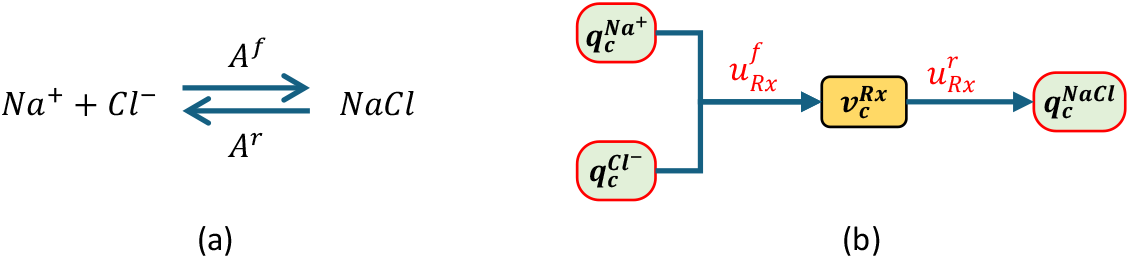
(a) A sodium-chloride reaction. (b) The bond graph model in simplified form.

The 0:node mass balance equations are:

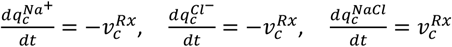

The reaction flux (inserting the Boltzmann equations for each chemical species) is

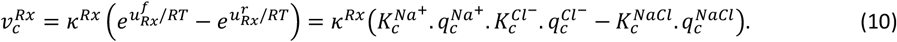

With 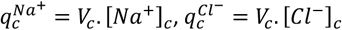, and 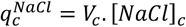 for compartment *c* of volume *V*_*c*_, equation (10) becomes

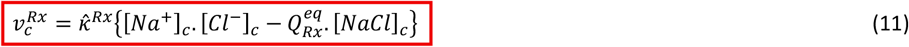

where 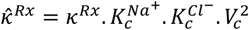 and 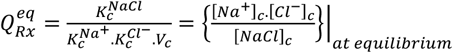.

This equation, now using concentrations, is the most useful form of the reaction flux, but it is important to remember that the ‘reaction rate’ constant 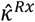 is now a combination of the kinetic parameter *κ*^*Rx*^ and the thermodynamic parameters 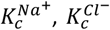. In practice 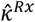 is determined experimentally by measuring the non-equilibrium flux, and the equilibrium constant 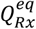 is determined by measuring the concentrations of Na^+^, Cl^-^ and NaCl at equilibrium.

#### (iii) Enzyme-catalysed reactions

An enzyme-catalysed reaction is illustrated in Figure 5.

**Figure 5.**
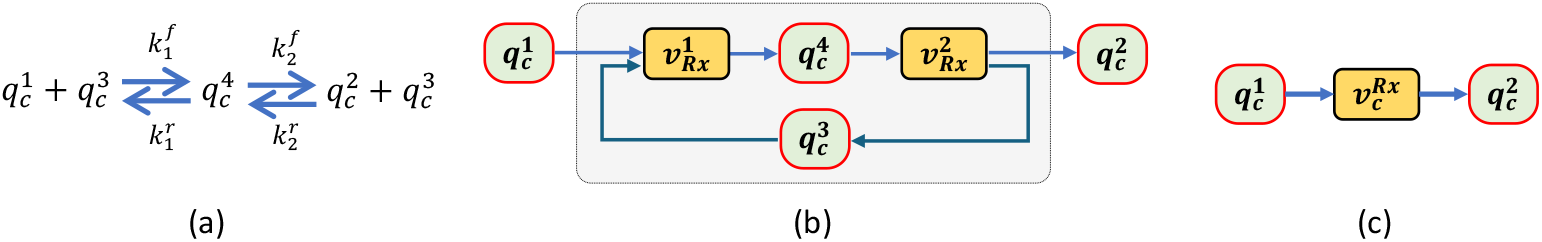
(a) An enzyme-catalysed reaction in which a solute 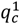 combines (reaction 1) with the enzyme 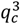 to form an intermediate 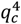 which then dissociates (reaction 2) to form the solute product 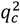 and recycled enzyme 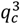. (b) The enzyme-catalysed reaction shown as a bond graph with the four red-bordered symbols representing both mass balance and storage, and the two reactions (with 1:nodes incorporated as in Figure 2) representing the points of energy balance (chemical stoichiometry). (b) A further simplification of the bond graph model in which the steady-state flux 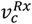 is the final net flux, computed from (19), for the whole reaction.

The flux balance equations, defined at the four 0:nodes, are

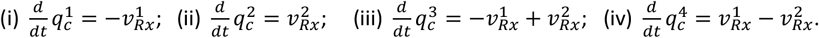

Note that, since 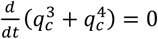, the total amount of enzyme is constant. i.e.,

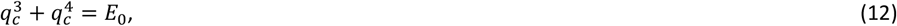

where *E*_0_ is the initial quantity of enzyme.

The energy balance equations, defined at the four 1:nodes, are

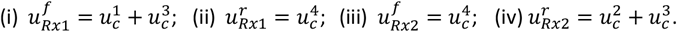

Using these potentials, the two reactions are

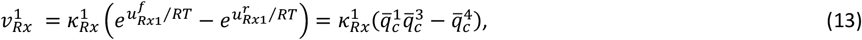

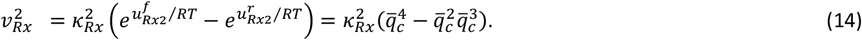

We can also express these fluxes in “mass action” form as

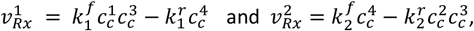

where 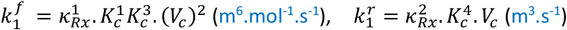,and 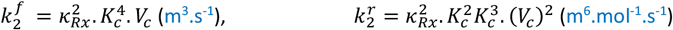.

Note the inconsistent units that result from combining reaction rate constants (mol.s^-1^) with thermodynamic constants (mol^-1^).

The *Briggs-Haldane assumption* [14] is that the amount of enzyme 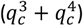 is much less than the amounts of substrate (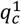 and 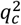), so that the unbound enzyme 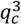 and the complex 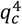 quickly reach a steady state (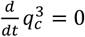 and 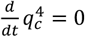). The second and third equations in (4) then give

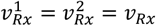

and hence, from (13) and (14),

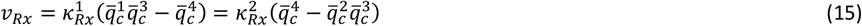

or

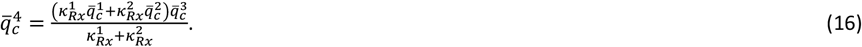

Also, from (12),

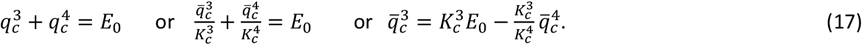

Substituting (17) into (16),

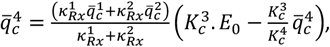

or

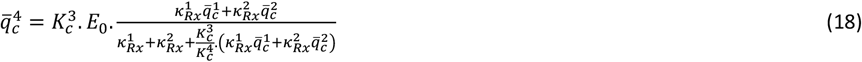

and hence, from (17),

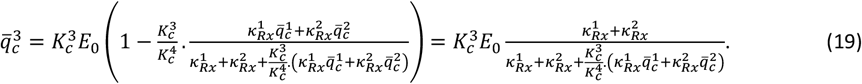

Substituting (18) and (19) back into the first equation in (15),

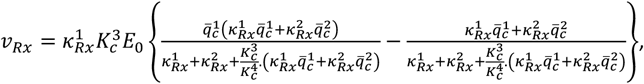

which simplifies to

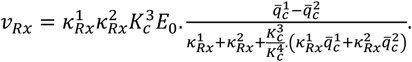

A convenient way of expressing this enzyme-catalysed reaction flux is

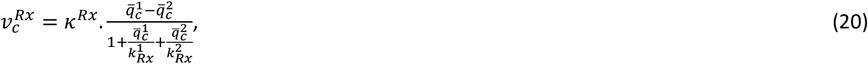

where the three parameters

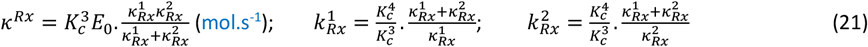

characterise the maximum flux, the value of 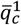 that corresponds to half that maximum when 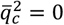. and the value of 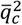 that corresponds to half that maximum when 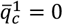. The flux term (20) shows saturation at high values of 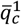 or 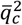. Note that we use 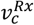 in (20) and in the bond graph diagrams in Figure 6, with *Rx* as a superscript to indicate that this is the steady-state flux for the whole enzyme-catalysed reaction. This convention greatly simplifies subsequent multi-enzyme reactions (e.g., for membrane transporters).

**Figure 6.**
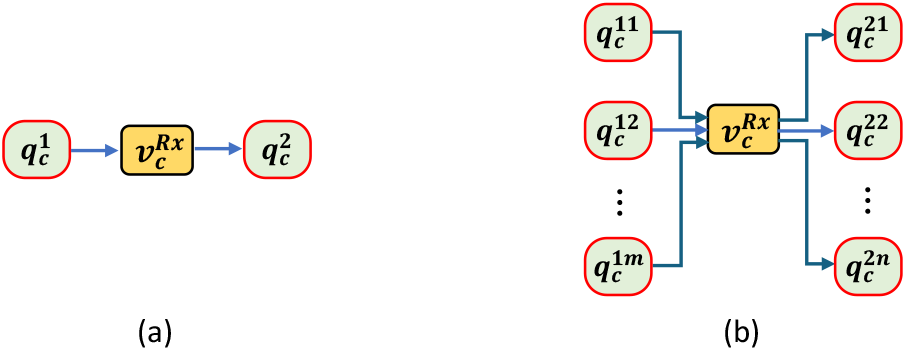
(a) The bond graph diagram for an enzyme-catalysed reaction under the Briggs-Haldane assumption where the flux term 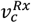 (given by equation (19)) now incorporates the action of the enzyme. (b) An enzyme-catalysed reaction with *m* reactants and *n* products.

If 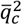 is assumed to be much smaller than 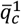 and is set to zero, equation (20) becomes the familiar Michaelis-Menten equation [14], but the reaction is then irreversible. The validity of the Briggs-Haldane assumption can be tested by comparing the solution of the full bond graph model with the reduced model given by (20).

Figure 6b shows the bond graph for an enzyme-catalysed reaction which has *m* reactants (assumed to bind simultaneously) and *n* products (assumed to unbind simultaneously).

Following the same process described above for the derivation of the full bond graph model, and then the corresponding reduced Briggs-Haldane model,

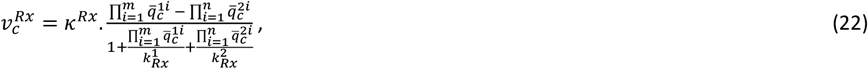

where the parameters 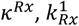 and 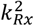, which characterise the catalysed reaction are given by (22).

The zero-flux state (*v*_*Rx*_ = 0) is reached when 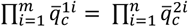.

The free energy of the reaction is

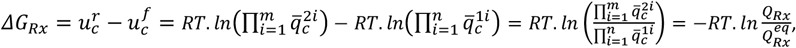

where

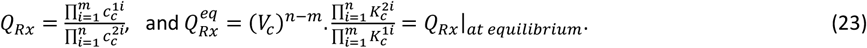

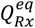 is the equilibrium constant for the reaction, which replaces the *m* + *n* thermodynamic coefficients.

The zero-flux state for the reaction corresponds to 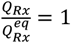 and *ΔG*_*Rx*_= 0.

Using (1) and (21), the flux expression (20) can be rewritten in terms of concentrations:

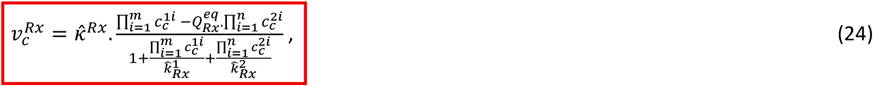

where

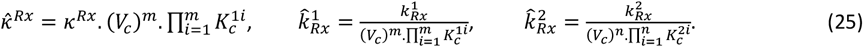

There are now four constants available 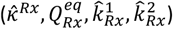 to fit the general enzyme catalysed reaction (24) to experimental data on the relationship between flux and the reactant and product concentrations.

For a very fast reaction (when the reaction rate constant *κ*^*Rx*^ → ∞), a finite (non-zero) value for the steady enzyme turn-over flux 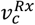 implies that the reaction is close to equilibrium 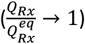.

#### (iv) ATP hydrolysis

ATP hydrolysis supplies the energy (usually via the ‘sodium battery’) to drive nearly all physiological processes [13]. The hydrolysis of ATP to ADP, P_i_ (inorganic phosphate), and H^+^ is represented chemically in Figure 7a and as an enzyme-catalysed bond graph process in Figure 7b.

**Figure 7.**
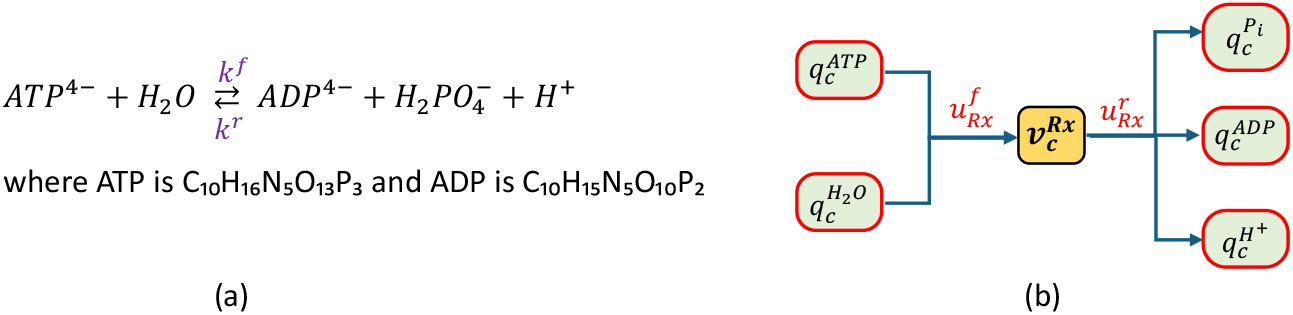
The bond graph representation of ATP hydrolysis. At pH 7, inorganic phosphate (*P*_*i*_) is a mixture of the protonated forms of phosphate: 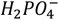 and 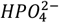. Here we assume 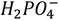 only.

This is an enzyme-catalysed reaction with an equilibrium constant, from (25), defined as

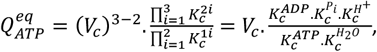

but determined from the experimentally measured concentrations at equilibrium:

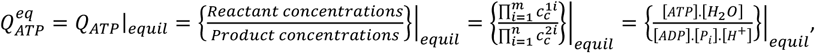

where the concentrations [..] are defined in units of mol.m^-3^, which is the same as millimol.L^-1^ or mM.

The concentration of water is 55.5 mM (the density 1 kg.L^-1^ divided by 0.018 kg.mol^-1^), but since the amount of water absorbed by the reaction is small compared to the quantity of water in which the reaction is taking place, a modified equilibrium constant is measured using

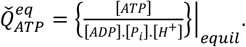

When measured at 298.15 K (25 °C) at pH 7 ([*H*^+^] = 10^-4^ mM and 1 mM [*Mg*^2+^] [18]), the equilibrium constant has the value 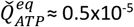 (assuming mM units) or, with *RT* = 2.5 kJ.mol^-1^,

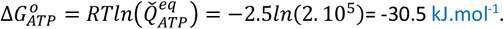

As we show below for the ATPase-driven NKA pump, in many situations ATP hydrolysis operates at Δ*G*s that are well in excess of this equilibrium.

#### (v) Glucose transporter (GLUT2)

Using a similar analysis (see [7] for details), the bond graph model of facilitated transport of glucose (‘Glc’) through the *SLC2A2*-encoded membrane transporter GLUT2 (see Figure 8), under the Briggs-Haldane assumption of steady state enzyme cycling and fast binding and unbinding, is given in terms of the non-dimensional intracellular and extracellular glucose amounts 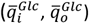 by

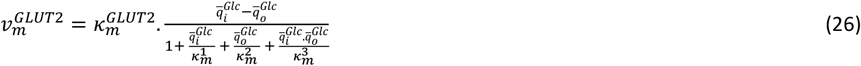

and, in terms of the intracellular and extracellular glucose concentrations ([*Glc*]_*i*,_ [*Glc*]_*o*_),

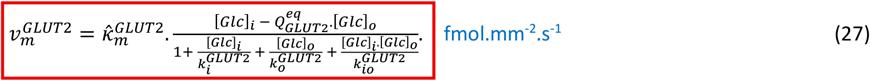

**Figure 8.**
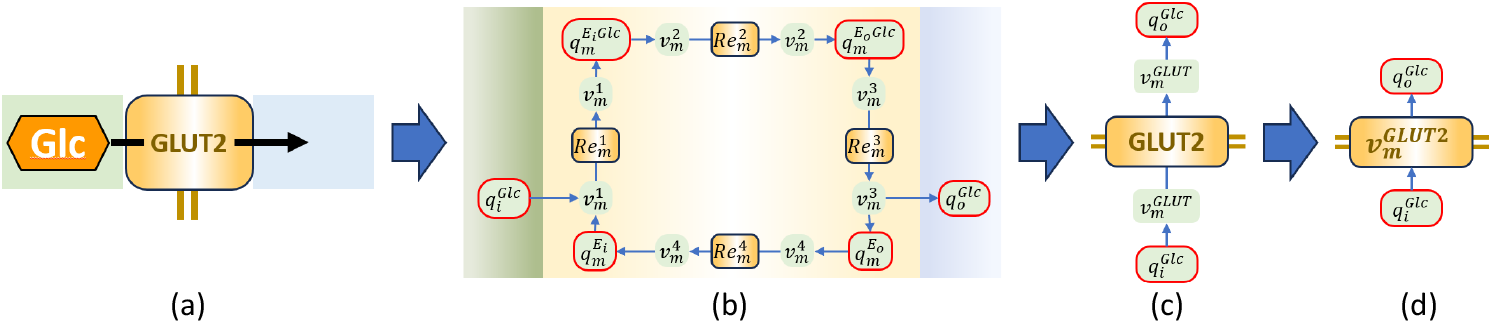
The GLUT2 transporter. (a) Facilitated transport of glucose across the cell membrane, shown by the double line (green background is inside the cell, blue outside); (b) Full bond graph model with reactions for 1. glucose binding, 2. translocation of the ligand-bound enzyme from inward-facing to outward-facing, 3. release of glucose, and 4. return of the enzyme to inward facing; (c) Reduced model where the flux 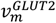 through the membrane is an algebraic function (equation 24) of the quantities of external and internal glucose; (d) the simpler representation in which the 1:nodes are included in the reaction.

For the analysis of cellular processes, we use units fmol (10^-15^ mol) for molar quantity, mm^2^ (10^-6^ m^2^) for area, and pL (10^-12^ L = 10^-15^ m^3^) for volume (1 fmol.pL^-1^ = 1 mol.m^-3^ = 1 mM). Since 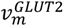 is a transmembrane flux, it is convenient to express as a molar flux per unit area of membrane, and therefore to use units of fmol.mm^-2^.s^-1^. Since the solute concentrations are in mM (mol.m^-3^), the reaction rate 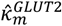 has units of fmol.mm^-2^.s^-1^/mol.m^-3^ or pL.mm^-2^.s^-1^ This parameter, fitted to whole cell data, is therefore a combination of reaction rate for the individual membrane protein transporter and a scaling constant that takes into account the areal protein density. When we later place this transporter into the whole cell model, we introduce an area-to-volume ratio for the cell that links concentrations and per-unit-area fluxes to changes in molar amounts.

Note that, from (7), the equilibrium constant is

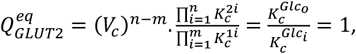

since the Boltzmann thermodynamic constant *K*_*c*_ is the same for internal and external glucose.

The parameters 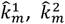 and 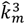 scale the relative contributions of extracellular and intracellular glucose and their product, respectively (see Table 1 for the value of these parameters fitted to data from Lowe and Walmsley [15]). 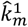 is the value of [*Glc*]_*i*_ that reduces the flux to 50% of its maximum value when [*Glc*]_*o*_ is zero, and 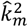 is the value of [*Glc*]_*o*_ that reduces the flux to 50% of its maximum value when [*Glc*]_*i*_ is zero. Note that 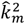 is an order of magnitude larger than 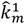 and that 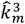 is a further order of magnitude larger than 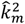. These parameters are expressed in terms of the thermodynamic and kinetic constants for the transporter as

**Table 1.** Parameter values used in the expression for flux 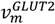 through the GLUT2 transporter. Values on the left are obtained by fitting (24) to data from Lowe and Walmsley [19]. Values on the right have been adjusted to reflect use of concentrations in (25).

| Parameters | Value | Unit | Parameters | Value | Unit |
| --- | --- | --- | --- | --- | --- |
| $K^{Glc}$ | ? | $\text{fmol}^{-1}$ | | | |
| $k_m^1$ | 1.474 | dimensionless | $k_i^{GLUT2}$ | 0.1094 | mM |
| $k_m^2$ | 21.67 | dimensionless | $k_o^{GLUT2}$ | 1.609 | mM |
| $k_m^3$ | 235.1 | dimensionless | $k_{io}^{GLUT2}$ | 1.296 | $(\text{mM})^2$ |
| $\kappa_m^{GLUT2}$ | 0.003284 | $\text{fmol.s}^{-1}$ | $Q_{GLUT2}^{eq}$ | 1 | dimensionless |
| | | | $\hat{\kappa}_m^{GLUT2}$ | 0.0442 | $\text{pL.s}^{-1}$ |

Figure 9 illustrates the flux through the steady-state (SS) bond graph model as a function of intracellular concentration, using equation (27) with the parameters given in Table 1.

**Figure 9.**
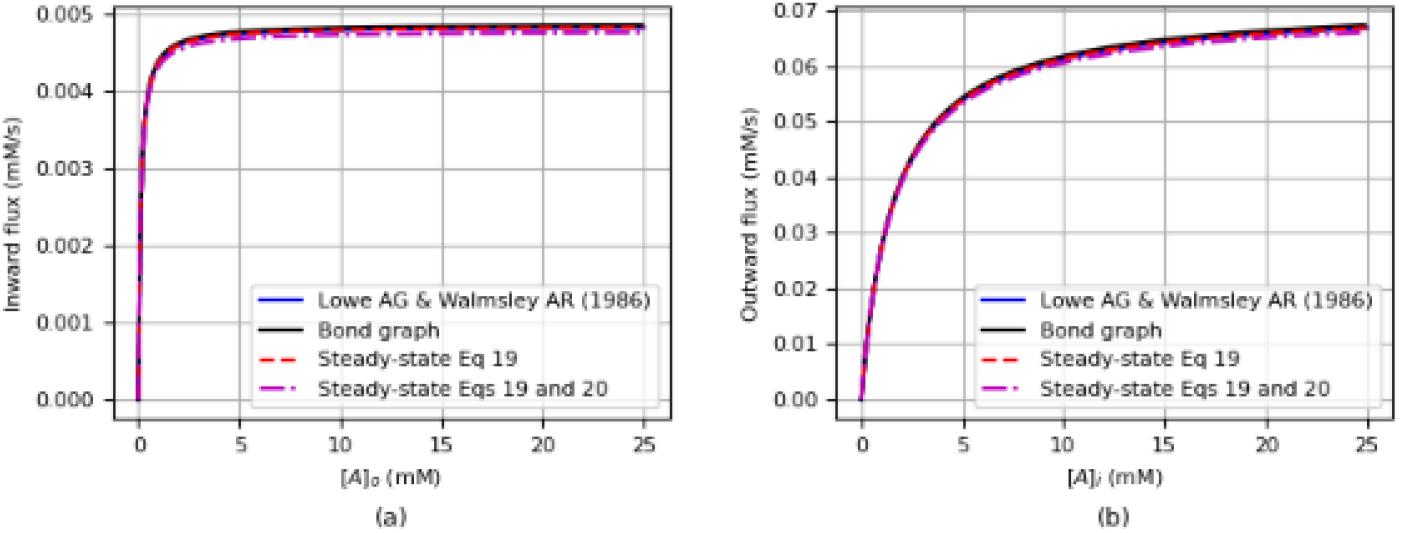
The GLUT2 bond graph model fitted to flux data from [19]. Reproduced from [7] with permission).

#### (vi) Electrogenic sodium-glucose cotransporter (SGLT1)

Glucose transport through the *SLC5A1*-encoded membrane co-transporter SGLT1 (see Figure 10), under the same assumptions used above (Briggs-Haldane and fast binding/unbinding) together with the assumption of no slippage (see [7]), is given in terms of non-dimensional quantities by

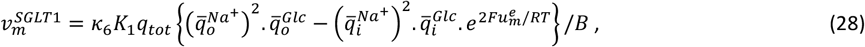

where

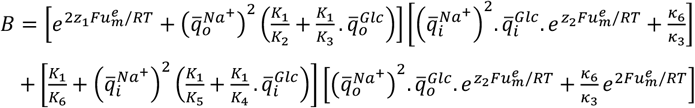

and *z*_1_ = *z*_2_ = 0.5.

**Figure 10.**
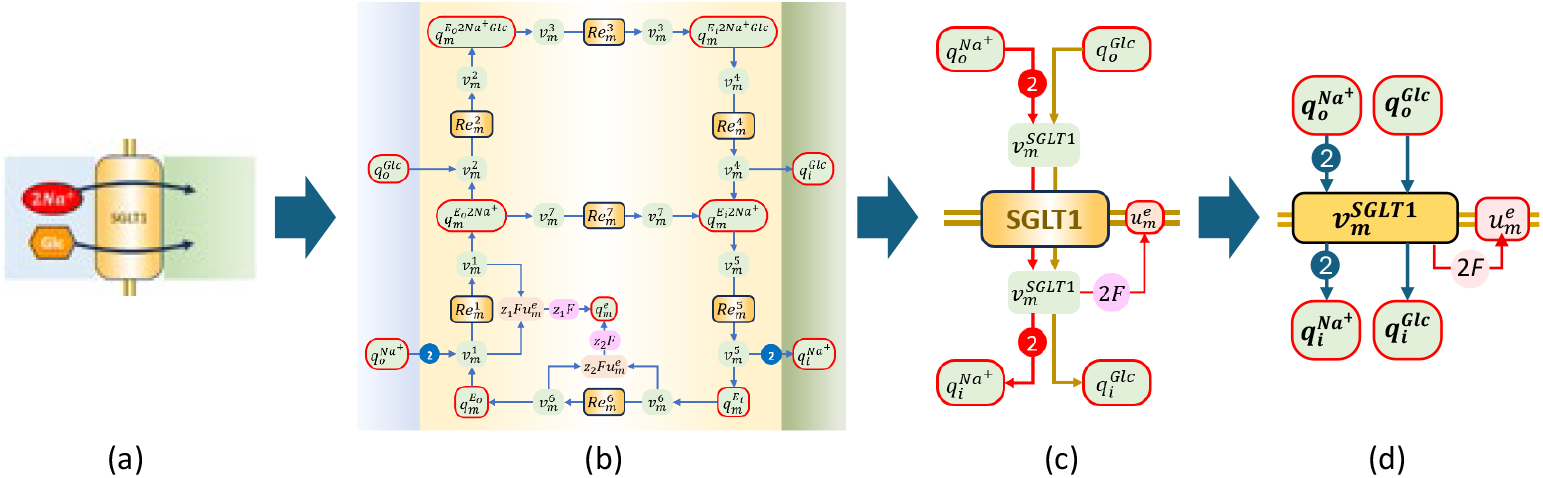
The sodium-glucose transporter SGLT1. (a) Symbolic representation for the transport of two moles of sodium for every one mole of glucose; (b) Full bond graph model; and (c) Reduced model where the flux 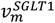 is an algebraic function (27) of the quantities of sodium and glucose either side of the membrane (see [7]); (d) the alternative representation in which the 1:nodes are included in the reaction.

Dividing numerator and denominator in (28) by 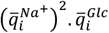, using concentrations rather than dimensionless molar quantities, and identifying the external compartment as ‘gut lumen’, (28) becomes

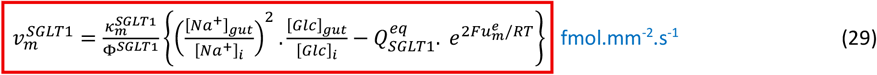

where

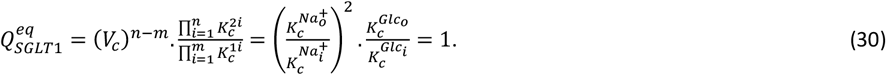

and the denominator term is

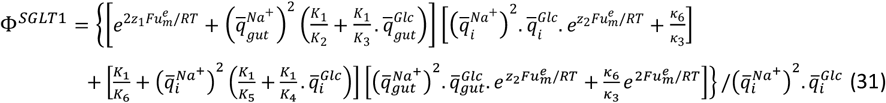

Note that 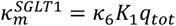 (fmol.mm^-2^.s^-1^) is a rate parameter also reflecting the overall expression level for the transporter and its reaction rate.

Figure 11 shows the flux through the steady-state (SS) bond graph model using the parameters given in Table 2.

**Table 2.** Parameter values used in the expression for flux 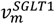 through the SGLT1 transporter. Values obtained by fitting (29) and (31) to data from Lowe and Walmsley [19] (see [7]}. The enzyme states corresponding to the thermodynamic parameters labelled *K*_1_.. *K*_6_, and the reactions corresponding to the kinetic parameters *κ*_3_ and *κ*_6_, are shown in the right-hand column.

| Parameters | Value | Unit | Solute & enzyme states |
| --- | --- | --- | --- |
| $K^{Na^+}$ | 0.0159 | fmol<sup>-1</sup> | $q_{gut}^{Na^+}, q_i^{Na^+}$ |
| $K^{Glc}$ | ? | fmol<sup>-1</sup> | $q_{gut}^{Glc}, q_i^{Glc}$ |
| $\kappa_m^{SGLT1}$ | 3.5e6 | fmol.mm<sup>-2</sup>.s<sup>-1</sup> | |
| $K_1$ | 3.66 | fmol<sup>-1</sup> | $q_m^{E_o}$ |
| $K_2$ | 330 | fmol<sup>-1</sup> | $q_m^{E_o 2Na^+}$ |
| $K_3$ | 321 | fmol<sup>-1</sup> | $q_m^{E_o 2Na^+ Glc}$ |
| $K_4$ | 321 | fmol<sup>-1</sup> | $q_m^{E_i 2Na^+ Glc}$ |
| $K_5$ | 905 | fmol<sup>-1</sup> | $q_m^{E_i 2Na^+}$ |
| $K_6$ | 0.314 | fmol<sup>-1</sup> | $q_m^{E_i}$ |
| $\kappa_3$ | 0.156 | fmol.s<sup>-1</sup> | $q_m^{E_o 2Na^+ Glc} \rightarrow q_m^{E_i 2Na^+ Glc}$ |
| $\kappa_6$ | 9.562 | fmol.s<sup>-1</sup> | $q_m^{E_i} \rightarrow q_m^{E_o}$ |

**Figure 11.**
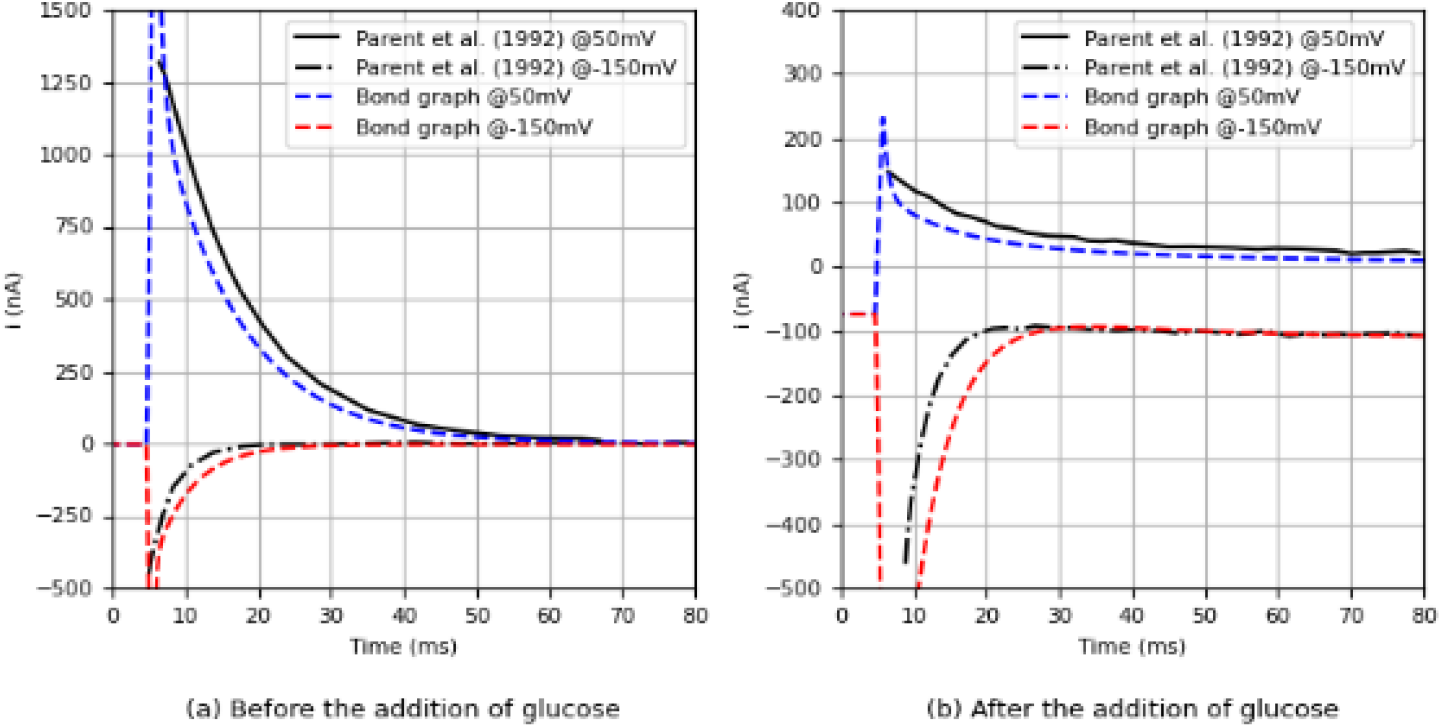
Shows SGLT1 model fitted to data (normalised flux). Reproduced from [7] with permission.

From (29) the reversal potential for SGLT1 (i.e. when 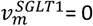) is given by

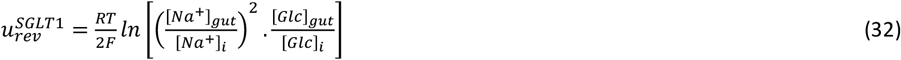

Substituting typical values (following a carbohydrate-rich meal) of [*Na*^+^]_*gut*_= 140 mM, [*Na*^+^]_*i*_= 15 mM, [*Glc*]_*gut*_= 40 mM, [*Glc*]_*i*_= 1 mM (with *RT*/*F*= 25 mV) gives 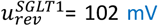. Given a typical epithelial cell membrane potential of about –50 mV, the SGLT1 cotransporter operates well away from the reversal potential. Note that at 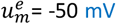, and with these sodium and glucose concentrations, the ratio of forward to reverse flux is ( 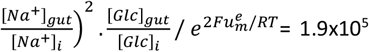, which means that 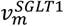 is not impeded by the rising [*Glc*]_*i*_ (which is also rapidly transferred to the blood by GLUT2).

#### (vii) Sodium-potassium ATPase pump (NKA)

The Na^+^-K^+^ ATPase NKA (a P-type ATPase) pumps 3 sodium ions out of the cell and 2 potassium ions in against their concentration gradients, using the energy supplied by the enzyme-catalysed hydrolysis of ATP [16]. The pump is electrogenic since there is a net movement of 1 charge for every ATP hydrolysed. By maintaining the sodium gradient across cell membranes, the NKA pump maintains the energy gradient used by many SLC transporters and is responsible for nearly 30% of the body’s resting metabolism. We have developed a 6-state model of NKA following the Post-Albers scheme [17], as illustrated in Figure 12 (see [9]).

**Figure 12.**
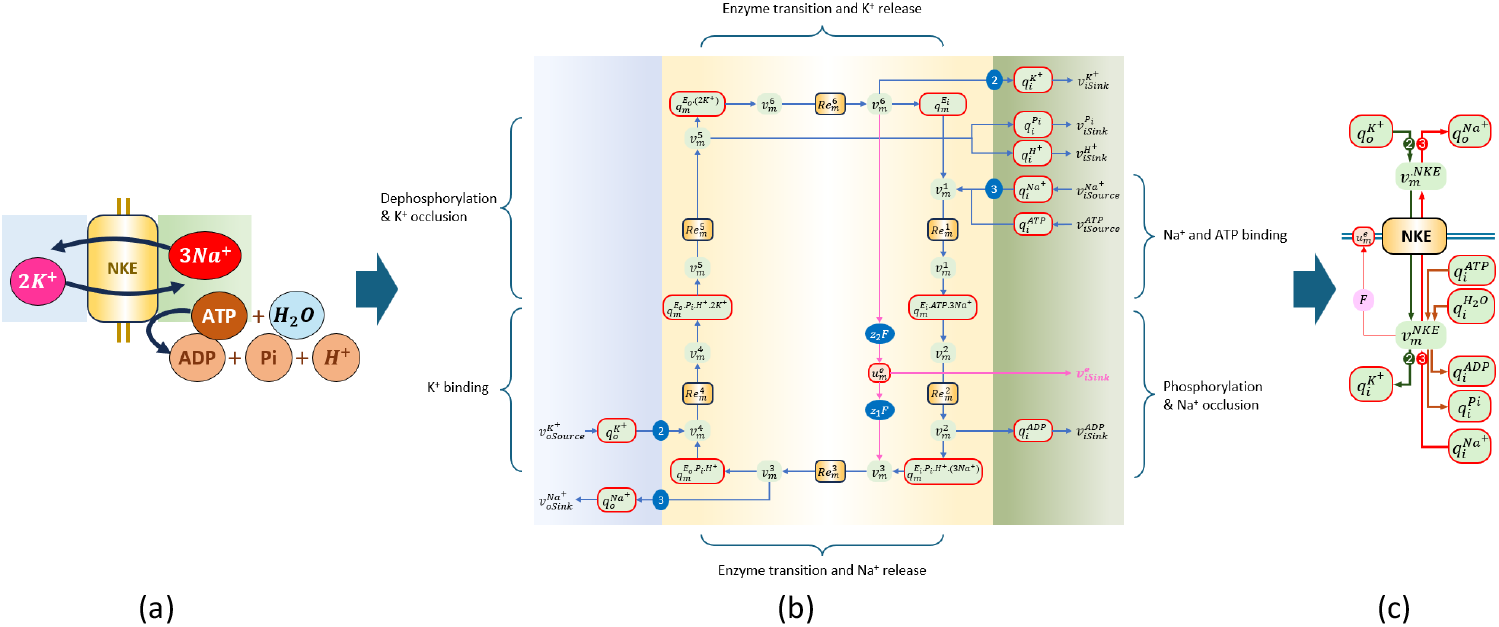
The sodium-potassium ATPase pump. (a) Symbolic representation for the transport of three moles of sodium outwards and two moles of potassium inwards for every mole of ATP hydrolysed; (b) The 6-state bond graph model; and (c) the reduced model for the flux 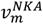.

The 6 states of the NKA protein, each followed by a reaction (defining the state transition), are (in clockwise order starting at the top right):

1. The first is the unbound inward-facing state 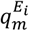 is followed by the membrane reaction 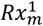 in which three intracellular sodium ions and one ATP molecule bind. Note that sodium binding is cooperative and that ATP binds to a catalytic site on the enzyme’s cytoplasmic domain with high affinity in the presence of bound sodium. The voltage sensitivity is minimal at this stage, as the binding of the three charged *Na*^+^s occurs on the cytoplasmic side without crossing the membrane’s electric field.
2. The second (inward facing) state 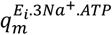 is followed by the 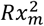 reaction in which bound ATP is hydrolysed and ADP is ejected. The other products of ATP hydrolysis, inorganic phosphate *P*_*i*_ and a proton *H*^+^, remain bound. The phosphorylation associated with ATP hydrolysis is thought to occlude the sodium ions within the membrane.
3. The third (inward facing) state is therefore 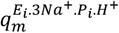 and is followed by reaction 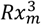 in which the protein transitions to outward facing and the three sodium ions are ejected (since the affinity for *Na*^+^ is greatly reduced by the inward-facing to outward-facing transition). This transition is highly voltage dependent as the three charged sodium ions are being transported outward against the membrane’s electric field gradient (and against a large *Na*^+^ concentration gradient). A more negative highly charged membrane (i.e., more negative membrane potential 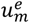) facilitates the reaction.
4. The fourth (now outward facing) state 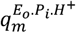 is followed by reaction 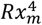, which binds two extracellular potassium ions. Note that cardiac glycosides, such as ouabain, bind to the 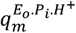 state, stabilizing it and blocking *K*^+^ binding.
5. The fifth (outward facing) state is therefore 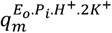. Potassium binding triggers dephosphorylation 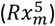 which occludes the bound *K*^+^ ions and ejects the remaining hydrolysis products (*P*_*i*_, *H*^+^). There is minimal voltage sensitivity.
6. The sixth and final (outward facing) state is 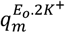. In the final reaction 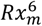, the two potassium ions are ejected into the intracellular space leaving the unbound protein ready to begin the next cycle. This *K*^+^ release involves moving 2 positive charges inward, facilitated by the membrane’s electric field gradient.

The expression for NKA flux [9] is

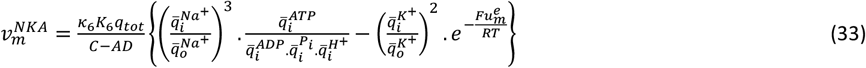

where the flux rate depends on the non-dimensional quantities *A, B, C, D* given in Table 4. See [9] for the meaning of each variable and parameter, and the derivation of these expressions. Model parameters are given in Table 3.

**Table 3.** Parameter values used in the expression for flux 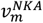 through the NKA transporter. Values obtained by fitting (34) to data from [18]. The enzyme states corresponding to the thermodynamic parameters labelled *K*_1_.. *K*_6_, and the reactions corresponding to the kinetic parameters *κ*_1_.. *κ*_6_, are shown in the right-hand column.

| Parameters | Value | Unit | Solute & enzyme states |
| --- | --- | --- | --- |
| $K^{K^+}$ | 0.0239 | fmol<sup>-1</sup> | $q_i^{K^+}, q_o^{K^+}$ |
| $K^{Na^+}$ | 0.0159 | fmol<sup>-1</sup> | $q_i^{Na^+}, q_o^{Na^+}$ |
| $K^{ATP}$ | 45.06 | fmol<sup>-1</sup> | $q_i^{ATP}$ |
| $K^{ADP}$ | 0.0015 | fmol<sup>-1</sup> | $q_i^{ADP}$ |
| $K^{P_i}$ | 0.8673 | fmol<sup>-1</sup> | $q_i^{P_i}$ |
| $K^{H^+}$ | 0.8673 | fmol<sup>-1</sup> | $q_i^H$ |
| $K_1$ | 0.01673 | fmol<sup>-1</sup> | $q_m^{E_i}$ |
| $K_2$ | 4.477e3 | fmol<sup>-1</sup> | $q_m^{E_i.ATP.3Na^+}$ |
| $K_3$ | 46.64e3 | fmol<sup>-1</sup> | $q_m^{E_i.P_i.H^+. (3Na^+)}$ |
| $K_4$ | 720.6 | fmol<sup>-1</sup> | $q_m^{E_o.P_i.H^+}$ |
| $K_5$ | 1.607 | fmol<sup>-1</sup> | $q_m^{E_o.P_i.H^+. 2K^+}$ |
| $K_6$ | 30.25e3 | fmol<sup>-1</sup> | $q_m^{E_o. (2K^+)}$ |
| $\kappa_1$ | 2.839e3 | fmol.s<sup>-1</sup> | $q_m^{E_i} \rightarrow q_m^{E_i.ATP.3Na^+}$ |
| $\kappa_2$ | 51.10 | fmol.s<sup>-1</sup> | $q_m^{E_i.ATP.3Na^+} \rightarrow q_m^{E_i.P_i.H^+. (3Na^+)}$ |
| $\kappa_3$ | 469.3 | fmol.s<sup>-1</sup> | $q_m^{E_i.P_i.H^+. (3Na^+)} \rightarrow q_m^{E_o.P_i.H^+}$ |
| $\kappa_4$ | 619.1e3 | fmol.s<sup>-1</sup> | $q_m^{E_o.P_i.H^+} \rightarrow q_m^{E_o.P_i.H^+. 2K^+}$ |
| $\kappa_5$ | 35.87 | fmol.s<sup>-1</sup> | $q_m^{E_o.P_i.H^+. 2K^+} \rightarrow q_m^{E_o. (2K^+)}$ |
| $\kappa_6$ | 868.4 | fmol.s<sup>-1</sup> | $q_m^{E_o. (2K^+)} \rightarrow q_m^{E_i}$ |
| $z_1$ | 0.9450 | dim | |
| $z_2$ | $z_1 - 1$ | dim | |

**Table 4.**
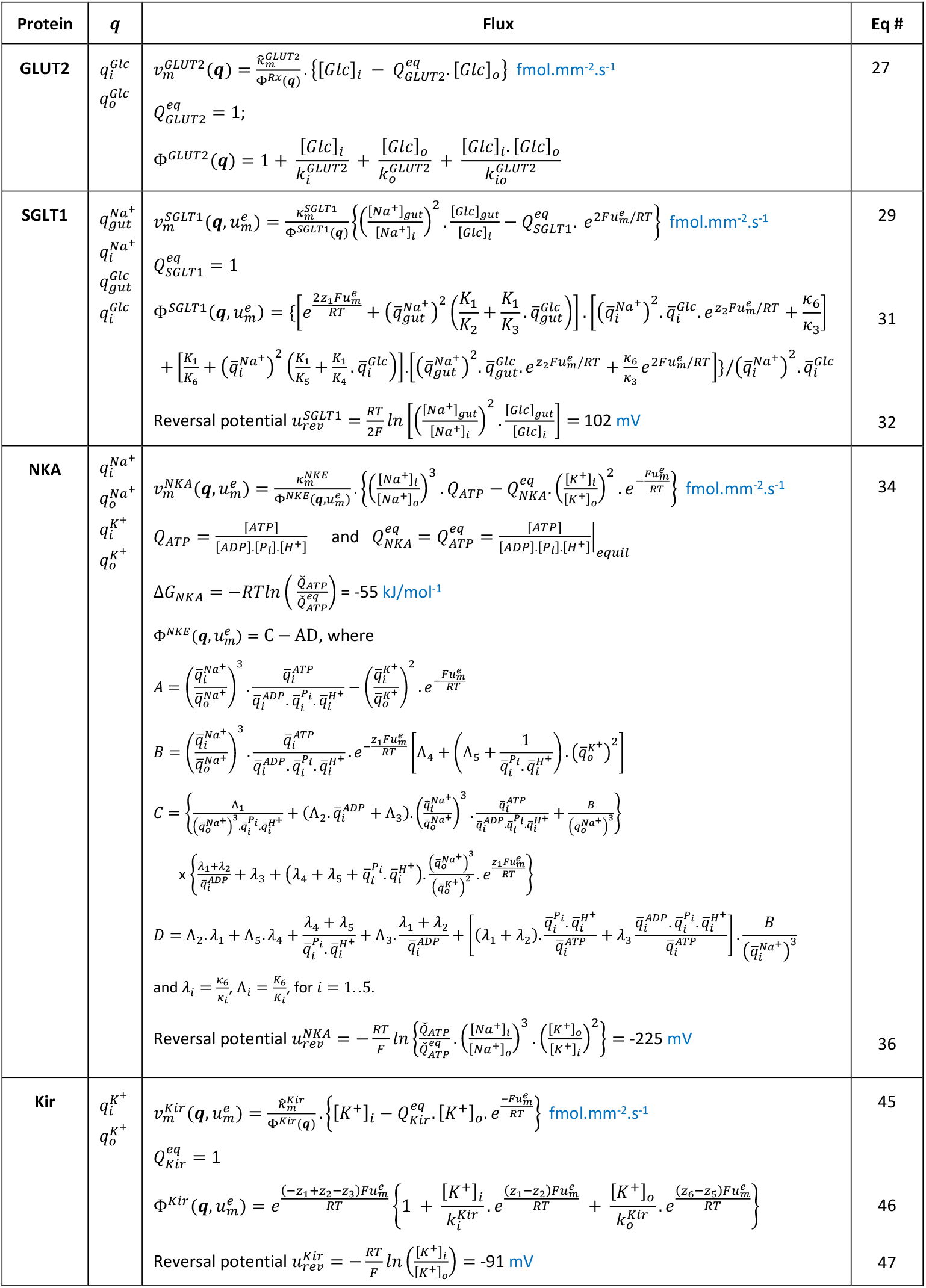
A list of the flux expressions in standardised form for the SLC transporters GLUT2 and SGLT1, the NKA pump, and the ion channel/transporter Kir. The reversal potentials shown for each membrane protein are calculated using values of [*Na*^+^]_*i*_= 15 mM, [*Na*^+^]_*o*_= 140 mM, [*K*^+^]_*i*_= 140 mM, [*K*^+^]_*o*_= 4.5 mM, [*Glc*]_*i*_= 1 mM, [*Glc*]_*o*_ = 40 mM, and 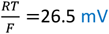.

Substituting for the concentrations using (1), equation (33) becomes

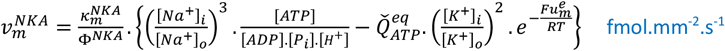

or

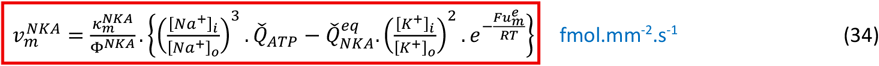

where 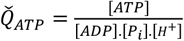 is the ratio of reactants to products for the intracellular components of ATP hydrolysis (assuming [*H*_2_*O*] constant) and 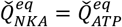 is the corresponding ratio of these components that attains equilibrium for ATP hydrolysis (i.e., the equilibrium constant for ATP hydrolysis). Note that other solute terms do not contribute to 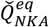 because the Boltzmann terms 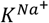 and 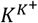 cancel out in the concentration ratio terms. The dimensionless ratio 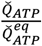 provides the active driving force for the reaction. 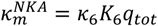 (fmol.mm^-2^.s^-1^) is a reaction rate constant, and Φ^*NKA*^ = *C* − *AD* is a nondimensional scaling factor for flux that is dependent on the concentrations of all solutes involved in the reaction (and is given by the expressions for *A, B, C*, & *D* in Table 4).

Figure 13 illustrates the dependence of flux (34) on 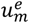 for varying values of [*Na*^+^]_*i*_ between 10 mM and 20 mM, with fixed values of [*Na*^+^]_*o*_= 140 mM, [*K*^+^]_*i*_= 145 mM, [*K*^+^]_*o*_ = 4.5 mM and a Gibbs free energy change of 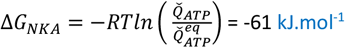.

**Figure 13.**
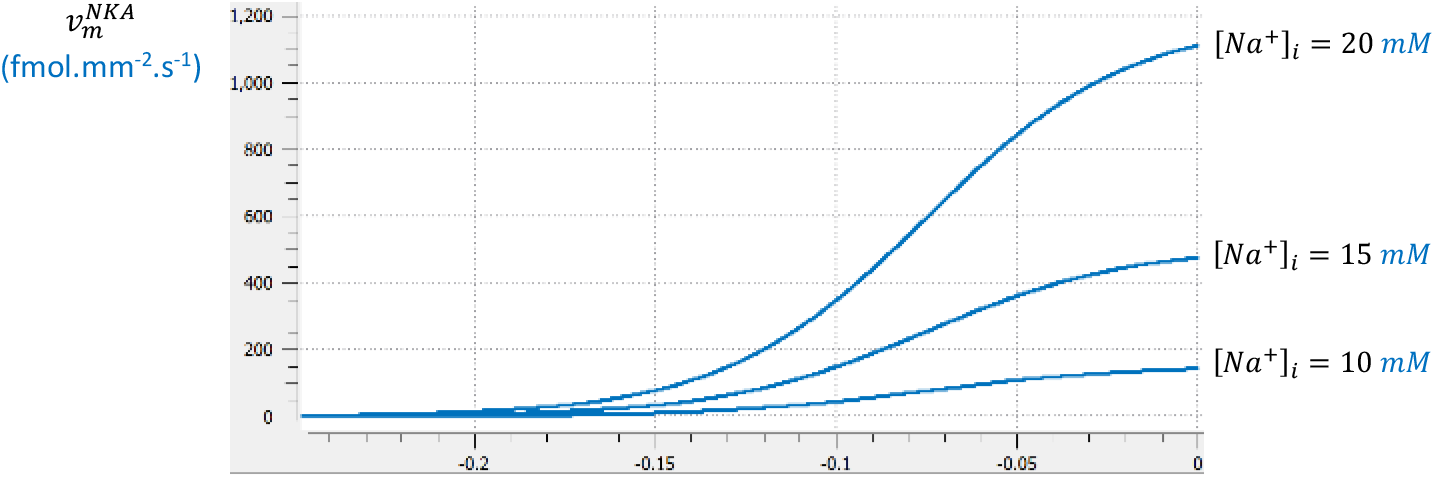
NKA transporter flux 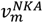 plotted against 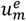 for three values of [*Na*^+^]^*i*^ (10 mM, 15 mM and 20 mM). Other concentrations are [*Na*^+^]_*o*_ = 140 mM, [*K*^+^]_*i*_ = 140 mM, [*K*^+^]_*o*_= 5 mM. The Gibbs free energy made available by ATP hydrolysis to drive the pump is 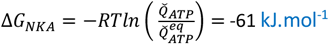. The reversal potentials corresponding to the intracellular sodium concentrations are (-238 mV, -271 mV, and -293 mV, respectively).

At equilibrium 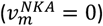, the ratio 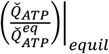, needed to maintain transmembrane ion gradients is

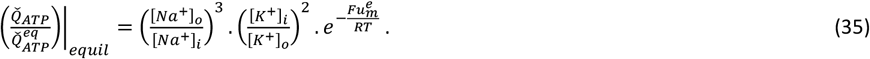

Applying 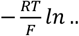 to both sides gives

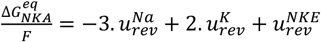

where 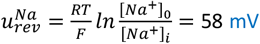 is the reversal (Nernst) potential for sodium (using 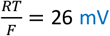), 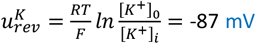 is the reversal (Nernst) potential for potassium, 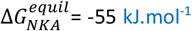 is the Gibbs free energy driving the equilibrium reaction, and 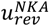 is the reversal potential for the NKA pump obtained from

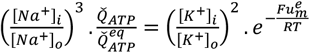

or

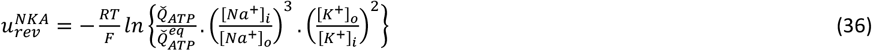

Substituting the standard concentrations into (36),

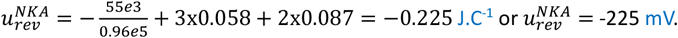

Note that a Gibbs free energy of –55 kJ.mol^-1^ corresponds to 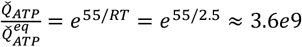, showing just how enormous the driving force is for maintaining intracellular sodium.

Equation (34) also illustrates the partitioning of Gibbs free energy:

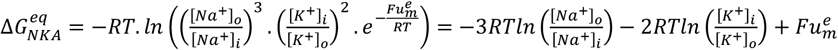

or

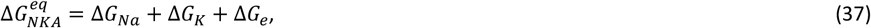

where 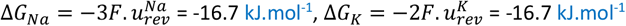 and Δ*G*_*e*_ = -21.6 kJ.mol^-1^ (which add to give 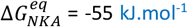 as above). Note that 39% (21.6/55) of the Gibbs free energy supplied by ATP hydrolysis at equilibrium goes into storing electrical energy in the cell membrane (by hyperpolarising it).

For this calculation of Δ*G*_*NKA*_ under equilibrium conditions, there is no heat loss because the NKA flux is zero; however, when the membrane voltage has the typically observed value of –50 mV, rather than being at the reversal potential of –225 mV, the free energy stored in the membrane capacitor drops to 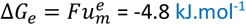, with the remaining 16.8 kJ.mol^-1^ dissipated as heat. The ‘efficiency’ of the pump is now 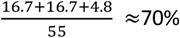, but of course the heat is helping to maintain body temperature, so it is more accurate to say that of the 55 kJ.mol^-1^ available from ATP hydrolysis, 61% goes to maintaining the electrolyte gradients, 9% goes to electrical energy storage in the membrane capacitance and 30% goes to thermal energy storage in the heat capacity of the cell and its surroundings.

#### (viii) The inwardly rectifying potassium ion channel (Kir)

We next consider the inwardly rectifying ion channel Kir (e.g., the Kir7.1 subtype). Highly selective, low-conductance single-ion channels can be considered as three- or four-state transporters (depending on the number of necessary intermediate steps). As illustrated in Figure 14, we use a three-state model with a bound state 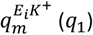, a meta-stable intermediate state 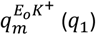, and an empty state 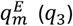. It is tempting to assume that the electrogenic step is associated with the transition between states 1 and 2 since this is where the ion transport occurs. But the bond graph model, while ensuring that physical conservation laws are satisfied for the protein as a whole, is limited in its ability to capture the details of the electrostatic field operating over the membrane and its interaction with the potassium ion moving through that field. The best we can do is to allow charge movement to be associated with all three reactions by including electrogenic effects via the unknown *z*_*i*_ (for *i*=1..6) terms as shown in Figure 14. The parameter estimation process described in Supplementary Material is used to determine these parameters (subject to the requirement that for each ion that crosses the membrane, the overall charge displacement for the membrane is -1).

**Figure 14.**
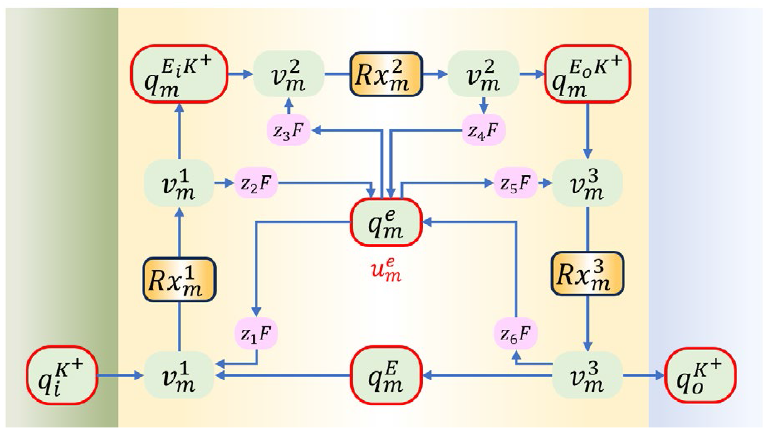
A 3-state bond graph model of the Kir inward rectifier with provision for charge movement by all reactions (represented by the purple transforming factors *z*_*i*_*F*). The binding and unbinding reactions (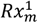 and 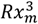) are assumed to be much faster than the protein transition from inward facing to outward facing 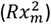.

From Figure 14, the flux equations associated with the three reactions are

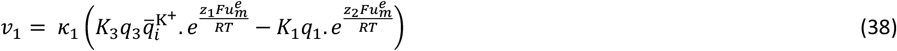

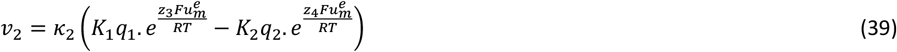

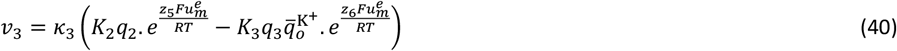

Assuming rapid binding and unbinding of K^+^ (*κ*_1_, *κ*_3_ → ∞), for finite values of *v*_1_ and *v*_3_, the bracketed terms on the right in (38) and (39) must be zero, and hence

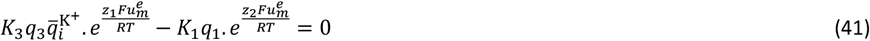

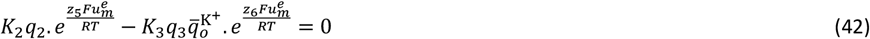

If we also assume (Briggs-Haldane) that the enzyme states are at steady-state, *v*_1_ = *v*_2_ = *v*_3_ = *v* and with substitutions for *K*_1_*q*_1_ and *K*_2_*q*_2_ from (41) and (42), respectively, (39) gives

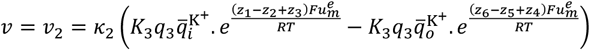

or

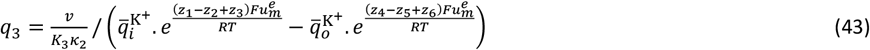

For a fixed (and specified) number of ion channels, *q*_*tot*_, and using (41) and (42),

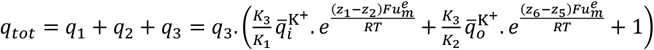

or, with (43),

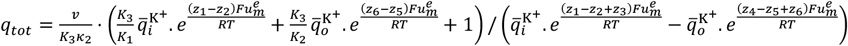

Rearranging for *v*,

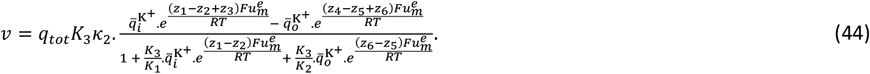

Finally, with 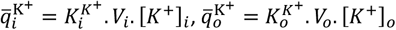, and defining 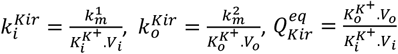 and 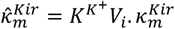, (44) becomes

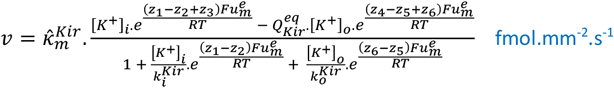

or, with −*z*_1_ + *z*_2_ − *z*_3_ + *z*_4_ − *z*_5_ + *z*_6_ = −1

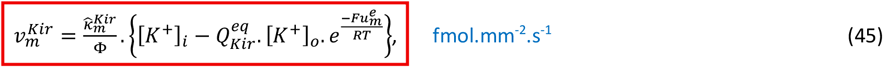

where

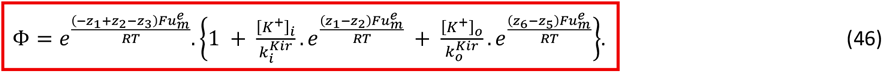

The electrical current is 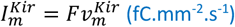 and the reversal or ‘Nernst’ potential is

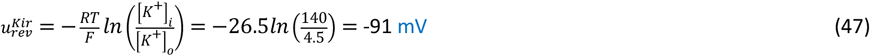

Figure 15 shows the current-voltage relation for the Kir channel.

**Figure 15.**
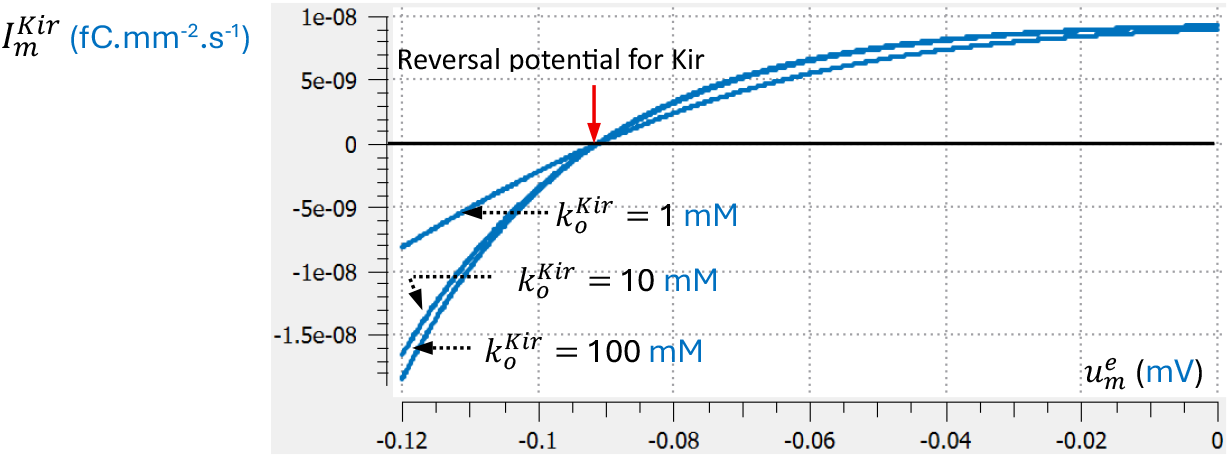
The current-voltage relation predicted for Kir by (45) and (46). Note the reversal potential at -91 mV and the inward rectification above this potential. Current-voltage curves are shown for 3 values of the parameter 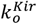. The curve steepens as this parameter is increased from 1 mM to 10 mM and then hardly changes for higher values. Changing the parameter 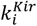 is equivalent to changes in 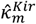 so this parameter is set to 1 mM.

The fluxes out of the intracellular solution and into the extracellular solution are

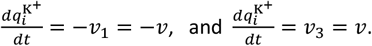

**Table 1.** Parameter values used in the expression for flux 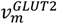 through the GLUT2 transporter. Values on the left are obtained by fitting (24) to data from Lowe and Walmsley [19]. Values on the right have been adjusted to reflect use of concentrations in (25).

| Parameters | Value | Unit |
| --- | --- | --- |
| $K^{K^+}$ | 0.0239 | $\text{fmol}^{-1}$ |
| $\hat{\kappa}_m^{Kir}$ | 0.0442 | $\text{pL}\cdot\text{s}^{-1}$ |
| $Q_{Kir}^{eq}$ | 1 | dimensionless |
| $k_i^{Kir}$ | 1 | mM |
| $k_o^{Kir}$ | 10 | mM |

### A COMMON FRAMEWORK FOR ALL TRANSPORTERS AND ATPase PUMPS

To provide a single unifying framework for all enzyme-catalysed reactions (including all membrane transporters, pumps and ion channels), we will express all steady-state reaction fluxes in the form:

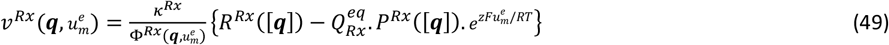

where ***q*** is a state vector that includes all solutes (with concentrations [***q***]) participating in the reaction, *R*^*Rx*^([***q***]) and *P*^*Rx*^([***q***]) are products of reactant and product concentrations, 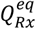 is the equilibrium constant given by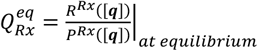, and 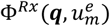 is an algebraic expression that affects the speed of *Rx* when the reaction is not at equilibrium. The constant *κ*^*Rx*^ scales the reaction flux and reflects the level of protein expression in a cell. Expressing all fluxes in this form clearly identifies:

1. the {}-bracketed component that governs equilibrium, including the equilibrium constant 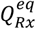,
2. the explicit dependence on membrane potential 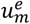 (for electrogenic reactions) that provides the reversal potential 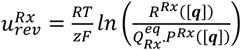,
3. the state-dependent term 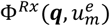 that governs the flux magnitude when not at equilibrium, the scaling constant *κ*^*Rx*^ that reflects the expression levels for the protein.

Table 4 summarises the transmembrane fluxes, all expressed in this form.

### A MODEL OF THE ENTEROCYTE

We now combine the GLUT2, SGLT1, NKA, and Kir models discussed above to create a model of homeostasis in the enterocyte, the absorptive cell of the epithelium in the small intestines. The NKA pump maintains a low intracellular [*Na*^+^]_*i*_ such that the SGLT1 cotransporter can use the sodium gradient between the gut lumen and the intracellular space to bring glucose into the cell. Glucose is metabolised into the ATP used to drive NKA and also flows out of the cell via GLUT2 facilitated diffusion. The NKA-driven potassium flux into the cell is balanced by potassium efflux through the Kir ion channel, as illustrated in Figure 16.

**Figure 16.**
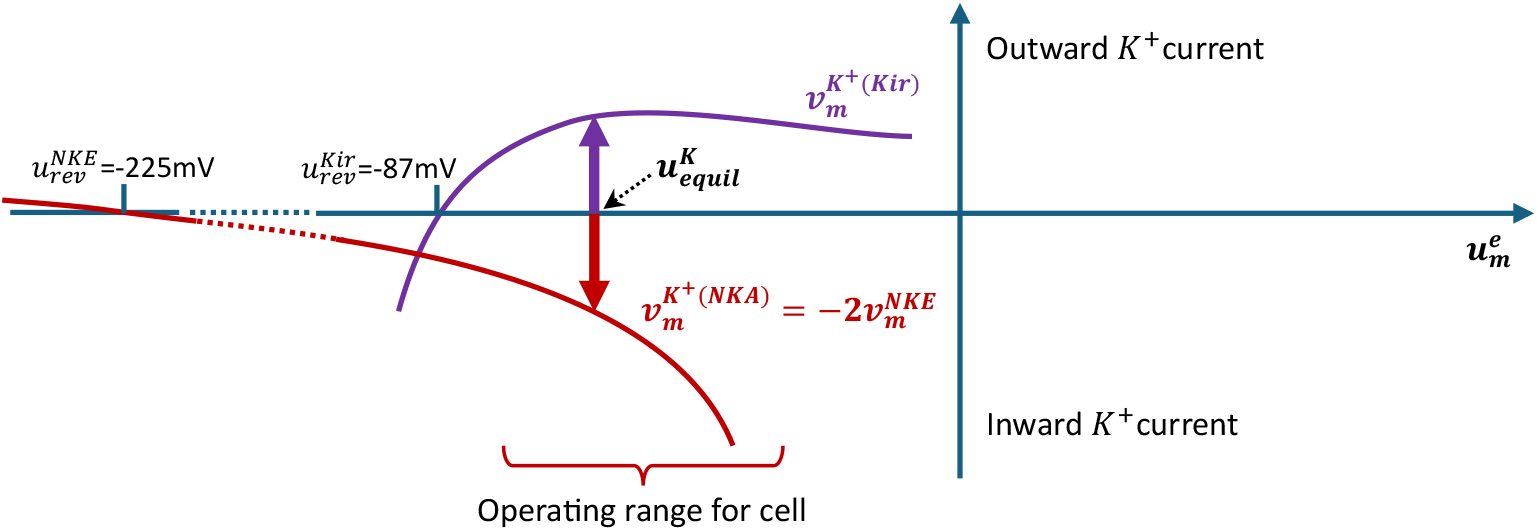
Equilibrium between the outward flow of *K*^+^ ions through Kir (when the membrane potential 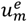 is above the reversal potential 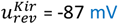) and the inward flow of *K*^+^ ions through NKA (when the membrane potential 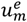 is above the reversal potential 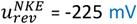 for NKA). Equilibrium is achieved when these two opposing currents are equal. The purple line shows the *K*^+^ ion efflux 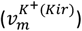 from Kir (equation [33]), and the dark brown line shows the influx 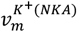 associated with NKA (equation (33) with two *K*^+^ ions transported for each NKA cycle). 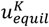 is membrane potential 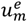 at this equilibrium.

**Figure 17.**
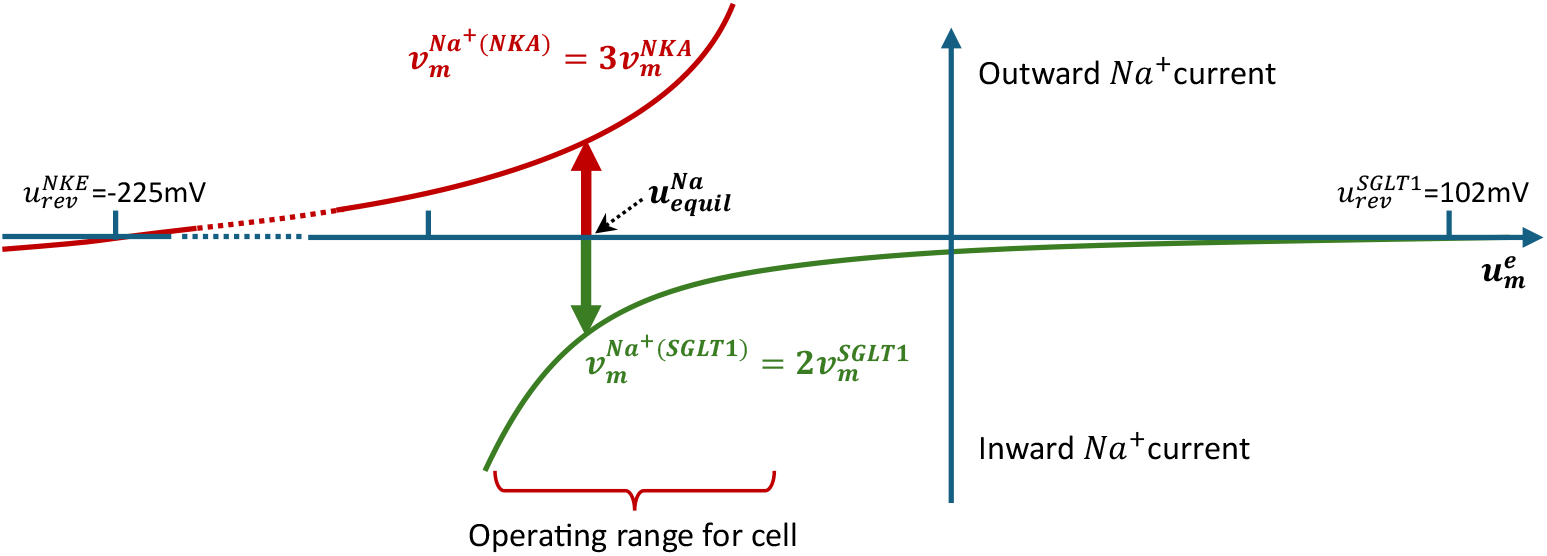
Equilibrium between the inward flow of *Na*^+^ ions through SGLT1 (when the membrane potential 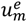 is well below the reversal potential 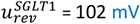) and the outward flow of *Na*^+^ ions through NKA (when the membrane potential 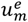 is above the reversal potential 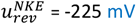 for NKA). Equilibrium is achieved when these two opposing currents are equal. The green line shows the *Na*^+^ ion influx 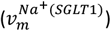 from SGLT1 (equation 29 with two *Na*^+^ ions transported for each SGLT1 cycle) and the dark brown line shows the efflux 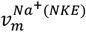 associated with NKA pump (equation (33) with three *Na*^+^ ions transported for each NKA cycle). 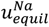 is the value of membrane potential 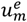 at this equilibrium.

Figure 16 illustrates the balance of inward sodium flux from SGLT1 with outward flux by NKA.

Figure 18 shows the various membrane processes and the glycolysis pathway that regenerates the ATP used by ATP hydrolysis in the NKA pump.

**Figure 18.**
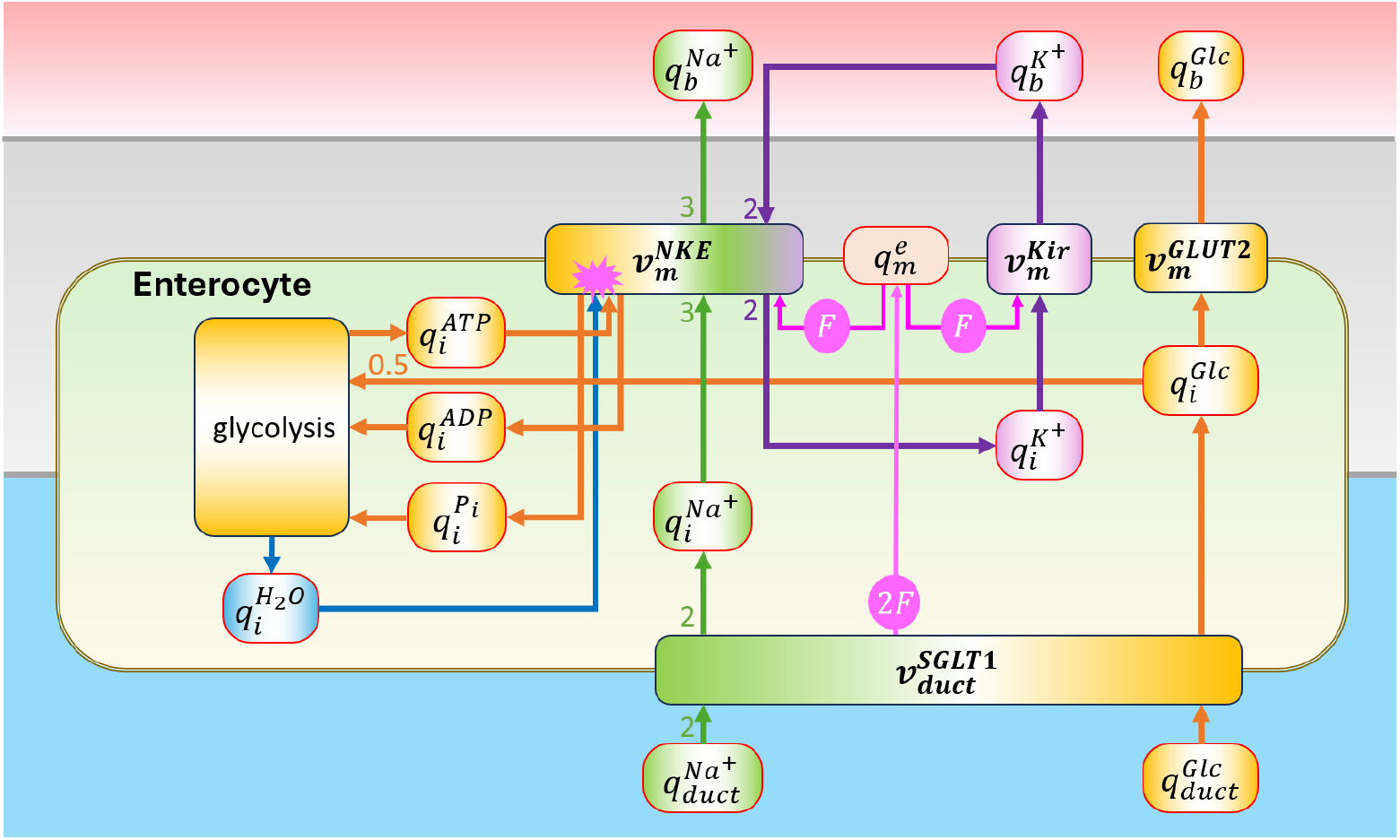
The enterocyte model includes NKA, Kir, and GLUT2 in the basolateral membrane and SGLT1 in the apical membrane. Energy is transferred between ***biochemical storage*** in ATP and solute concentrations, ***electrical storage*** in membrane capacitance, ***mechanical storage*** in the cell wall compliance (which determines the pressure-volume behaviour associated with water movement), and ***heat storage*** in the thermal capacity of water in the cell and its surroundings. One mole of glucose is assumed to instantaneously convert 2 moles of ADP to ATP via glycolysis with the generation of 2 moles of water.

The membrane voltage is 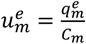 is calculated from net influx of membrane charge:

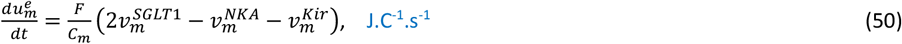

where *C*_*m*_ is the membrane capacitance (assumed to be 1 μF.cm^-2^ or 10^-8^ C^2^.J^-1^.mm^-2^). Note that

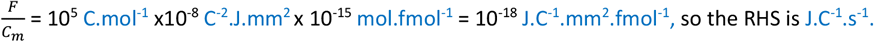

Increases in intracellular potassium and sodium ion concentrations are given by

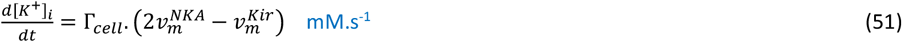

and

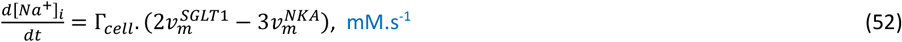

where Γ_*cell*_ (m^-1^) is the ratio of membrane area A_*cell*_ to cell volume V_*cell*_. Note that we are assuming the same areal density of the NKA pump, Kir channel, and GLUT2 transporter (all in the basolateral membrane in contact with interstitial fluid and effectively with blood) and the SGLT2 cotransporter (in the apical membrane in hence in contact with the intestinal lumen).

Since 1 mol of glucose is assumed to replace 2 mol of ADP with 2 mol ATP instantaneously, the intracellular glucose is given by

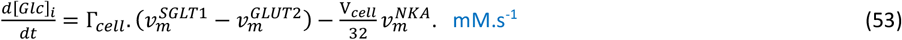

To examine how these four membrane proteins achieve homeostasis of sodium, potassium and glucose in the enterocyte, we first consider the two transporters Kir and NKA. Figure 19(a) shows the reaction fluxes 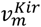 and 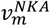 as a function of membrane potential 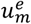 from -120 mV to 0 mV under physiologically normal conditions ([*Na*^+^]_*i*_= 10 mM, [*Na*^+^]_*o*_= 140 mM, [*K*^+^]_*i*_= 140 mM, [*K*^+^]_*o*_= 4.5 mM and 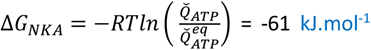). The reversal potentials for the two membrane proteins are 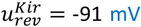 and 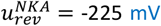, respectively. The effect on *K*^+^ flux of these two transporters is shown in Figure 19(b) where the net *K*^+^ flux 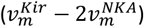 is plotted against 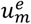 for different values of [*Na*^+^]_*i*_ from 0 mM to 12 mM. For [*Na*^+^]_*i*_ = 0 mM the net flux is zero at -91 mV and then occurs at progressively less negative membrane potentials as [*Na*^+^]_*i*_ increases, up to a limit of little over 10 mM. Beyond this there is no balance possible because the influx of *K*^+^ by NKA (associated with the high efflux of *Na*^+^) exceeds the efflux of *K*^+^ achieved by Kir at all potentials.

**Figure 19.**
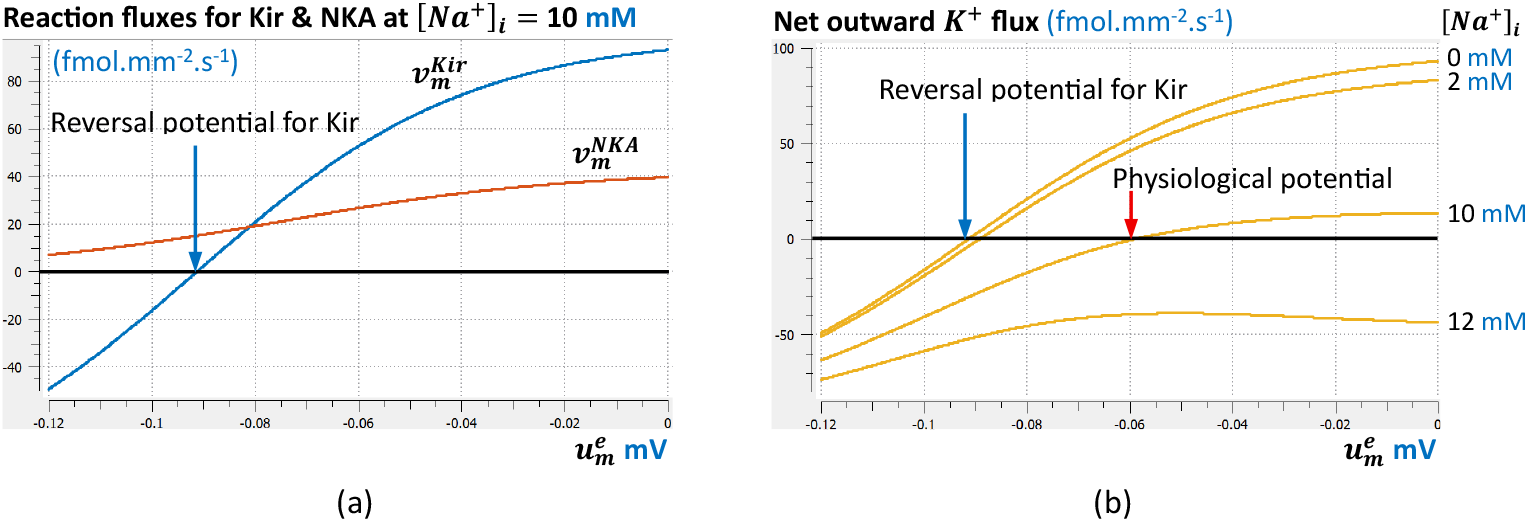
(a) Fluxes 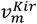 and 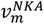 through Kir and NKA as a function of membrane potential 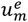. The reversal potentials for Kir and NKA under standard physiological conditions are -91 mV and -225 mV, respectively. (b) The net outward potassium flux 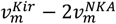 shown as a function of 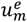 for values of [*Na*^+^]_*i*_ from 0 mM to 12 mM. For the specified electrolyte concentrations ([*Na*^+^]_*o*_= 140 mM, [*K*^+^]_*i*_= 140 mM, [*K*^+^]_*o*_= 4.5 mM) there is a well defined zero net *K*^+^ flux for values of [*Na*^+^]_*i*_ from 0 mM to 10 mM, but not for values of [*Na*^+^]_*i*_ above 10 mM.

The flux balance for *Na*^+^ depends on both NKA and SGLT1. Figure 20 shows how the balance of sodium efflux 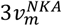 and sodium influx 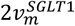 (i.e. a net efflux of 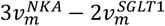) depends on membrane potential 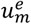.

**Figure 20.**
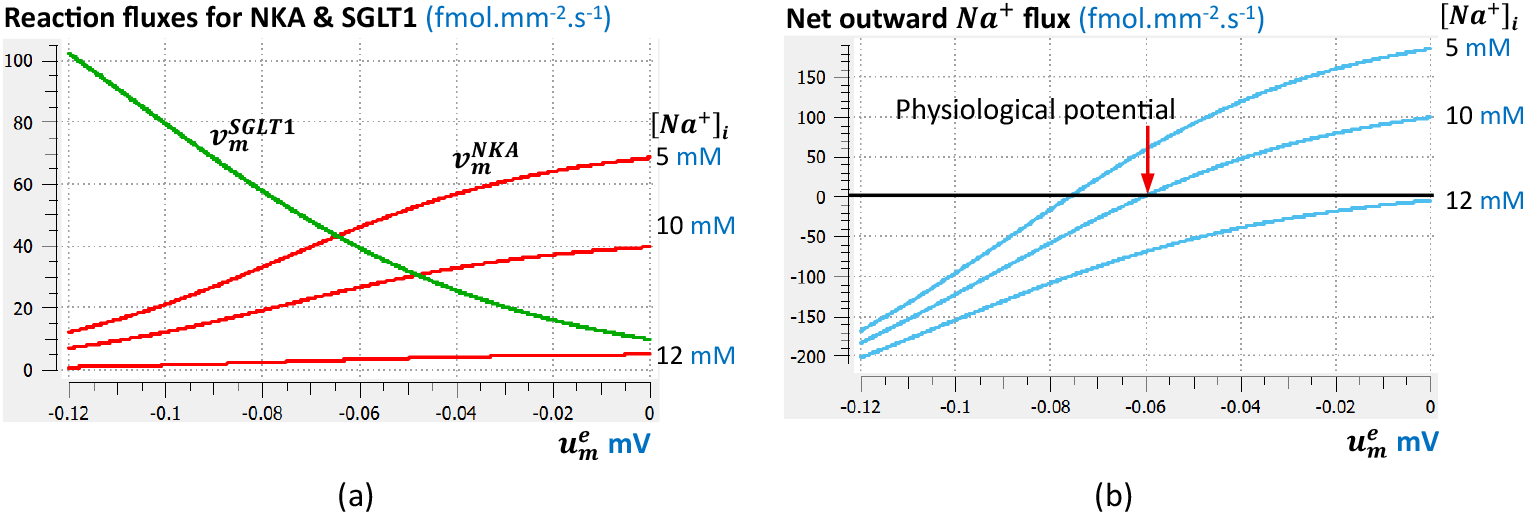
(a) Fluxes 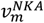 and 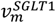 through NKA and SGLT1 as a function of membrane potential 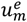. Both fluxes are computed for 3 values of [*Na*^+^]_*i*_, but this has little effect on 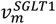. (b) The corresponding net outward *Na*^+^ flux (calculated from 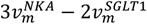).

Achieving zero net flux for both [*Na*^+^]_*i*_ and [*K*^+^]_*i*_ also ensures zero net flow of charge across the membrane since all charge transfer is associated with those ions.

At [*Na*^+^]_*i*_= 10 mM, both the net *K*^+^ flux and the net *Na*^+^ flux are zero at the same potential 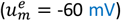. These values of [*Na*^+^]_*i*_ and 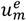 were calculated from (50) and (51) for a fixed values of 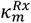 for each of the three transporters. An alternative is to set the left hand sides of (50) and (51) to zero for all values of 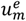 and compute the corresponding scaling on the 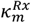 values (as a function of 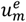) in order to achieve zero net flux. The computed scale factors are shown as a function of 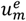 in the bottom panel of Figure 21. The unscaled fluxes (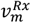 for each of the transporters) are shown in the top panel.

**Figure 21.**
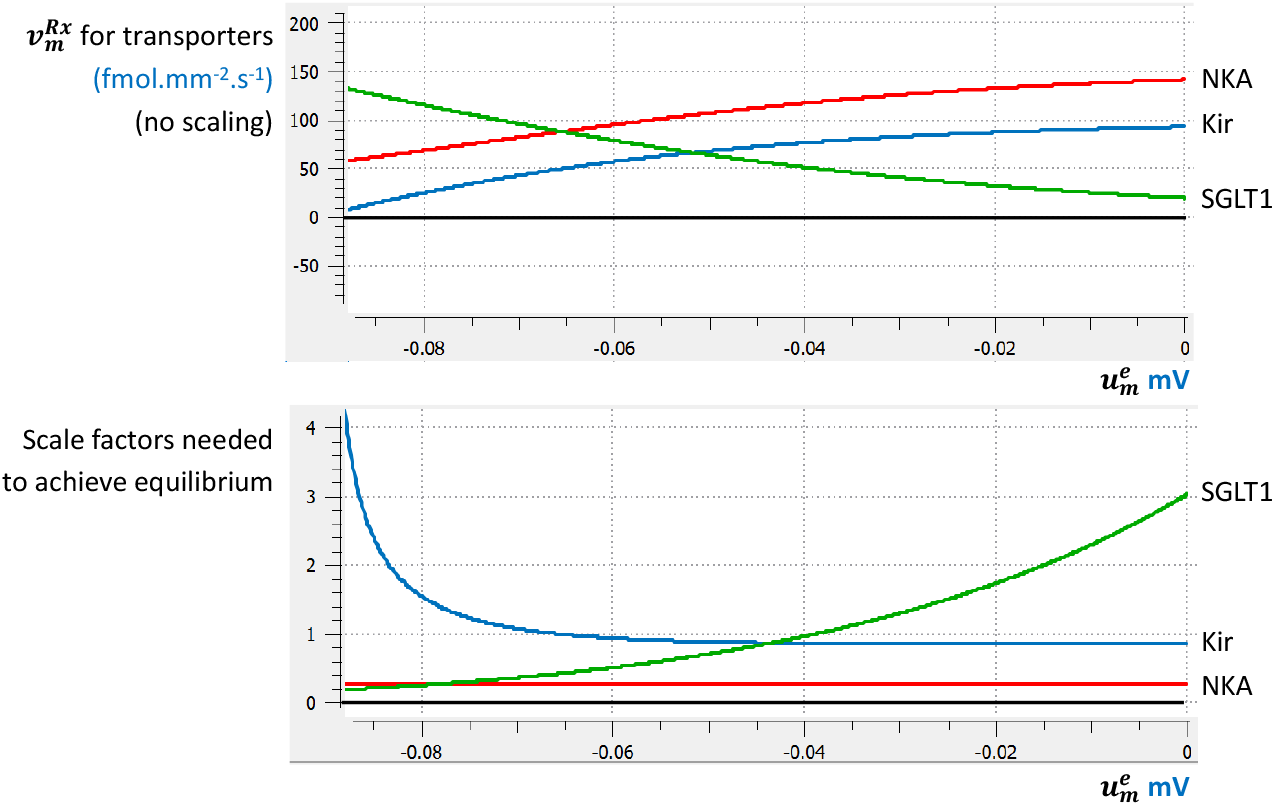
The transporter fluxes (top) and scaling on the reaction rate constants (bottom) needed for Kir and SGLT1 (relative to that on NKA) to achieve homeostasis of [*Na*^+^]_*i*_ and [*K*^+^]_*i*_. These relative scaling values depend on the membrane potential 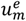 at which the net fluxes 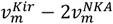 and 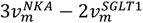 are both zero (i.e. these net fluxes are zero for all values of 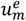 shown. Note the rapid increase in scaling on 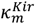 in the bottom graph as 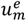 approaches the reversal potential for Kir (i.e. 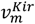 approaches zero on top graph).

The relative scale factors for SGLT1 and Kir shown in the bottom panel of Figure 21 would correspond to either regulation of the protein kinetics (e.g., by phosphorylation) or to the protein expression levels (under transcriptional control).

## DISCUSSION

Epithelial cells lining the small intestines take up glucose and sodium (delivered via ingested food) via the SGLT1 transporter, using solute gradients to drive the transmembrane flux (two sodium ions for each glucose molecule). Intracellular glucose is immediately converted to ATP via glycolysis (rather than oxidative metabolism), which then drives the NKA pump to maintain an intracellular sodium concentration of about 10 mM in the face of 140 mM sodium in the capillaries. The 140 mM/10 mM sodium difference provides a ‘battery’ that drives many other transmembrane transport processes. Note that a ‘battery’ is a source of potential energy and only loses energy when current flows (sodium ions in this case). The ‘sodium battery’ (i.e., the Gibbs free energy available from the sodium gradient) is therefore maintained by the Gibbs free energy of ATP hydrolysis – essentially the ‘top-up battery’ needed to maintain the sodium battery.

In this paper, we have explained how the physical principles of conservation of mass, conservation of charge, and conservation of energy are conveniently implemented in models of biological processes using bond graphs. The bond graph approach is well-known in the engineering literature for problems that involve energy transfer between the four available forms of energy storage – chemical, electrical, mechanical, and thermal. Physiological processes almost always involve energy transmission, exchange, and conversion (including dissipation) between all four physical domains, and bond graphs therefore provide an ideal framework for modelling these processes. We show how an appropriate choice of a small number of units (joules, meters, seconds, moles, coulombs, and entropy or Kelvin) provides a common foundation for describing all physical processes, and we illustrate the application of bond graphs to basic biochemical processes where Gibbs free energy is key to understanding and modelling reactions.

We then derived biophysically based bond graph models of GLUT2, SGLT1, Kir, and the NKA ATPase pump and coupled NKA with the transport of glucose via SGLT1 and GLUT2. Glycolysis supplies the ATP needed by NKA for maintaining homeostasis of [*Na*^+^]_*i*_ and [*K*^+^]_*i*_.

Given the 3:2 ratio of sodium efflux to potassium entry in NKA, the 2:1 ratio of coupled sodium to glucose entry by SGLT1, and the single potassium efflux by Kir, the cycling rates of SGLT1 and Kir must be 1.5x and 2x that of NKA, respectively, in order to maintain homeostasis of [*Na*^+^]_*i*_ and [*K*^+^]_*i*_, and hence also of the membrane resting potential 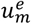. This balance of the transporter fluxes can either be achieved by adjusting the levels of [*Na*^+^]_*i*_[*K*^+^]_*i*_, and 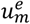, or by altering either the phosphorylation status or expression levels of the transporters, as illustrated in Figure 21. Future work will explore the regulation of these membrane transport proteins.

